# *Cauliflower mosaic virus* protein P6 forms a microenvironment for RNA granule proteins and interferes with stress granule responses

**DOI:** 10.1101/2022.10.26.513884

**Authors:** Gesa Hoffmann, Silvia Lopéz-Gonzaléz, Amir Mahboubi, Johannes Hanson, Anders Hafrén

**Affiliations:** Department of Plant Biology, Uppsala BioCenter, Swedish University of Agricultural Sciences, 75007 Uppsala, Sweden; Linnean Center for Plant Biology, 75007 Uppsala, Sweden; Umeå Plant Science Centre, Department of Plant Physiology, Umeå University, Umeå, Sweden

**Author notes:** The author(s) responsible for distribution of materials integral to the findings presented in this article in accordance with the policy described in the Instructions for Authors (www.plantcell.org) is: Anders Hafrén.

## Abstract

Biomolecular condensation is a multipurpose cellular process that viruses use ubiquitously in their multiplication. CaMV replication complexes are condensates that differ from most viruses in being non-membranous assemblies and consist of RNA and protein, mainly viral protein P6. Despite description of these viral factories already half a century ago with many observations that followed since, functional details of the condensation process, their properties and relevance has remained enigmatic. Our main findings include a large dynamic mobility range of host proteins within viral factories, while the viral matrix protein P6 is immobile in accordance with representing the central node of these condensates. As novel components of VFs we identify stress granule (SG) nucleating factors G3BP7 and the UBP1 family. Similarly, as SG components localize in VFs during infection, ectopic P6 localizes to SGs and reduces their assembly after stress. Intriguingly, it appears that soluble rather than condensed P6 suppresses SGs and mediates also other essential P6 functions, suggesting that the increased condensation over the infection time-course may accompany a progressive shift in selected P6 functions. Together, this study highlights VFs as dynamic condensates and P6 as a complex modulator of SG responses.

## Introduction

The post-transcriptional fate of mRNA is controlled by an extensive network that mediate RNA processing, translation and ultimately degradation (Chantarachot and Bailey-Serres, 2018). An intriguing feature is the compartmentalization of many of these processes in membrane-less condensates termed RNA granules, including nucleoli, cajal bodies and paraspeckles in the nucleus and, processing bodies (PBs) and stress granules (SGs) in the cytoplasm (Spector, 2006). Studies on the physical properties of RNA granules suggest that they assemble through liquid-liquid phase separation, a process influenced by RNA-RNA interactions, as well as RNA interactions with protein low-complexity or prion-like domains (Uversky, 2017; Alberti et al., 2019; Youn et al., 2019; Boncella et al., 2020). RNA granules can assemble in a matter of minutes upon stress induction and disperse just as quickly when translationally favorable conditions are re-entered (Weber et al., 2008). This makes them a highly adaptable environment that offers possibilities for rapid and diverse molecular crowding and compartmentalization.

The most extensively studied cytoplasmic RNA granules in plants and other organisms are SGs and PBs, both implicated in the transient storage of non-translating RNA (Kedersha and Anderson, 2007; Decker and Parker, 2012; Chantarachot and Bailey-Serres, 2018; Guzikowski et al., 2019). An anticorrelation between ribosome- and SG-associated mRNAs was established in animals and plants (Sorenson and Bailey-Serres, 2014; Khong et al., 2017). Despite an enormous progress in the field, the functional importance of the actual assembly into microscopically visible RNA granules for mRNA regulation is still largely unclear (Guzikowski et al., 2019). Both, SGs and PBs have the capacity to influence mRNA translation, storage, and decay, as well as protein signaling and also serve as protective refuges during stress. They are induced by RNA release from polysomes triggered by abiotic or biotic stresses (Riggs et al., 2020). Despite representing distinct structures, they also share protein components and appear to even fully overlap in specific conditions (Buchan et al., 2013; Youn et al., 2018; Frydryskova et al., 2020), supporting a dynamic interface. In the “mRNA cycle” model (Buchan and Parker, 2009), mRNA movement between SGs, PBs and ribosomes is considered to be both dynamic and bidirectional. Notably, the efficiency by which RNAs enter SGs is highly variable but correlates positively with RNA length and poor translation (Khong et al., 2017). Viral RNAs can be extremely long and polycistronic, together with antiviral translational responses, making them especially prone to SG regulation. Indeed, there are numerous examples of animal viruses being targeted by SGs (Poblete-Duran et al., 2016), and these RNA granules would appear to be an integral part of the hosts antiviral defense, as well as a major target for manipulation by viral effectors (Lloyd, 2016; Poblete-Duran et al., 2016; Miras et al., 2017; Pooggin and Ryabova, 2018; Jaafar and Kieft, 2019; Stern-Ginossar et al., 2019). The extent to which plant virus infections involve the SG pathway remains largely elusive. UBP1, represented by the RBP45/RBP47/UBP1 family in Arabidopsis, is the closest homologue of the mammalian SG nucleation factor TIA-1 (Lorkovic and Barta, 2002; Weber et al., 2008; Gutierrez-Beltran et al., 2021), and was identified to assemble in non-canonical RNA granules and suppress translation of Potato virus A (Hafren et al., 2015). Another report suggested that two unrelated plant viral proteins can suppress SGs through their interactions with G3BP (Krapp et al., 2017), another core SG nucleating factor (Abulfaraj et al., 2018; Yang et al., 2020). Assuming a similar co-dependence of plant viruses as observed for animal viruses, there is a large unknown to uncover for the green viruses interacting with SGs.

CaMV is a double-stranded DNA pararetrovirus that replicates in large amorphous cytoplasmic inclusion bodies referred to as viral factories (VFs). These VFs have been described as non-membranous electron-dense protein/RNA-rich structures alike RNA granules. Despite that VFs are considered functionally associated with virus genome replication, translation and particle assembly (Schoelz and Leisner, 2017), our current understanding of these eminent inclusions and their dynamic functions is limited. The main component of VFs is the viral protein P6. P6 interacts with all other CaMV proteins (Himmelbach et al., 1996; Hapiak et al., 2008; Lutz et al., 2012), as well as a multitude of host proteins to exert it’s many functions. One of the key functions of P6 is to regulate viral translation involving physical interactions with ribosomal proteins, namely L13 (Bureau et al., 2004), L18 (Leh et al., 2000) and L24 (Park et al., 2001) and translational regulators eIF3g (Park et al., 2001) and TOR (Schepetilnikov et al., 2011). However, the spatial relation between VFs and P6 in association with the translation machinery is still vague but could be at the surface of VFs as these were found decorated by ribosomes (Shepherd, 1976). We are interested in the dynamics and functions of CaMV VF associated components and found recently that the PB components DCP5, LSM1a and VCS localize in VFs and support virus translation (Hoffmann et al., 2022). This raised a question of analogy between RNA granules and VF matrixes, which we pursued in the current study that identified and characterized SG components as novel factors in VFs.

## RESULTS

### P6 forms immobile matrixes of viral factories and self-condensates

The viral factories (VFs) are essential structures in CaMV infection and are largely build by P6, but the aggregation dynamics and P6 mobility within VFs are unknown. We monitored the VF size and shape maturation along the course of infection using the transgenic marker lines 35S:P6-GFP and 35S:P6-mRFP. In the absence of infection, P6-GFP formed numerous foci spanning a broad size range similar to what has been observed previously (Harries et al., 2009), while P6-mRFP was mainly soluble and formed much fewer, smaller and more uniform sized foci (Figure 1A-C). When P6 marker lines where infected with CaMV, both P6-GFP and P6-mRFP re-localized to mark the characteristic large amorphous VFs formed during infection (Figure 1A). A quantification showed that the number of foci decreased while the size and irregularity increased between 14- and 21-days post infection (dpi) (Figure 1B-D). This can be explained by fusion of smaller P6 foci to eventually mature into few but large VFs per cell, in accordance with previous interpretations (Shepherd, 1976).

**Figure 1:**
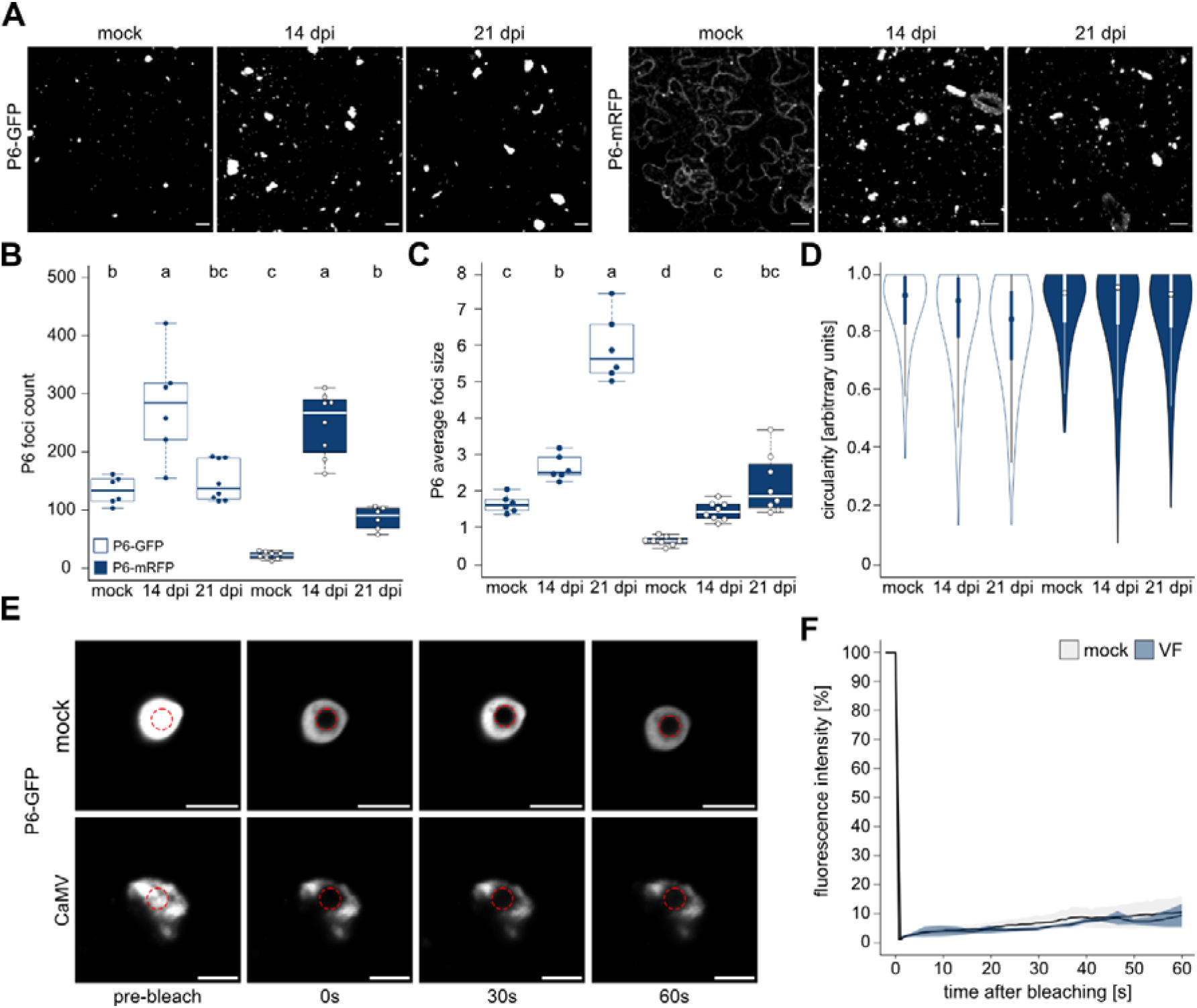
P6 establishes an immobile matrix in viral factories. **(A)** Maturation of P6 foci during infection time course (noninfected, 14 and 21 dpi) with CaMV was followed in P6-GFP and P6-mRFP marker lines by microscopy. Representative images are confocal Z-stack projections (Scale bars = 10 μm). **(B)** The amount of P6-GFP and -mRFP foci in 100 × 100 μm^2^ at timepoints corresponding to (A). Counts were obtained with a custom ImageJ pipeline using six to eight replicate images. **(C)** Average size of P6 foci in stacks corresponding to (B). Values were calculated from six to eight replicates with a custom ImageJ pipeline. **(D)** Circularity distribution of P6 foci at each time point as determined by ImageJ circularity masking. **(E)** FRAP analysis of P6-GFP in mock conditions (n=7) and in VFs at 21 dpi (n=8). Photobleached region is indicated by a red circle. Scale bars = 5 μm. **(F)** Normalized fluorescence intensities in FRAP analysis corresponding to (E) are plotted against time after bleaching. Solid lines represent mean, shades denote ± standard deviation. Letters in B-C indicate statistical groups determined by one-way ANOVA followed by Tuckey’s HSD test (α = 0.05).

While having a markedly different degree of condensation in uninfected conditions, P6-GFP and P6-mRFP behaved analogously by abundant accumulation in VFs during infection. To estimate the mobility of P6 in these structures, we used fluorescence recovery after photobleaching (FRAP) analysis on P6-GFP and P6-mRFP tagged VFs from CaMV infection as well as P6-GFP condensates in the absence of infection (Figure 1E and F, supplemental Figure 1). P6 was near static with very little recovery observed in all conditions, supporting that P6 is largely immobile in VFs and forms a robust VF matrix.

### Core SG components localize to VFs

We recently showed that PB components localize in VFs during CaMV infection (Hoffmann et al., 2022) and opted to further our understanding of the nature of these condensates, in particular to what extent they resemble cytoplasmic mRNA granules. To evaluate SG components in VFs, we established marker-lines of canonical SG proteins, namely GFP-RBP45c, GFP-RBP47b, GFP-RBP47c, GFP-UBP1b, GFP-UBP1c and GFP-G3BP7 in Col-0. The six markers localized in a diffuse nucleocytoplasmic pattern and assembled into microscopical foci upon heat stress (supplemental Figure 2A). In CaMV infected tissue, however, all markers localized to the large amorphous foci characteristic of VFs in addition to similar sized SG-like foci observed in response to heat (Figure 2A). By using a double marker line expressing GFP-RBP47b and P6-mRFP, we further found by a co-localization analysis that GFP-RBP47b indeed localizes to VFs with almost all P6 signal being co-marked by GFP-RBP47b (Figure 2B and C, “M1”) and about 40% of the total GFP-RBP47b signal being present in P6 foci (Figure 2B and C, “M2”).

**Figure 2:**
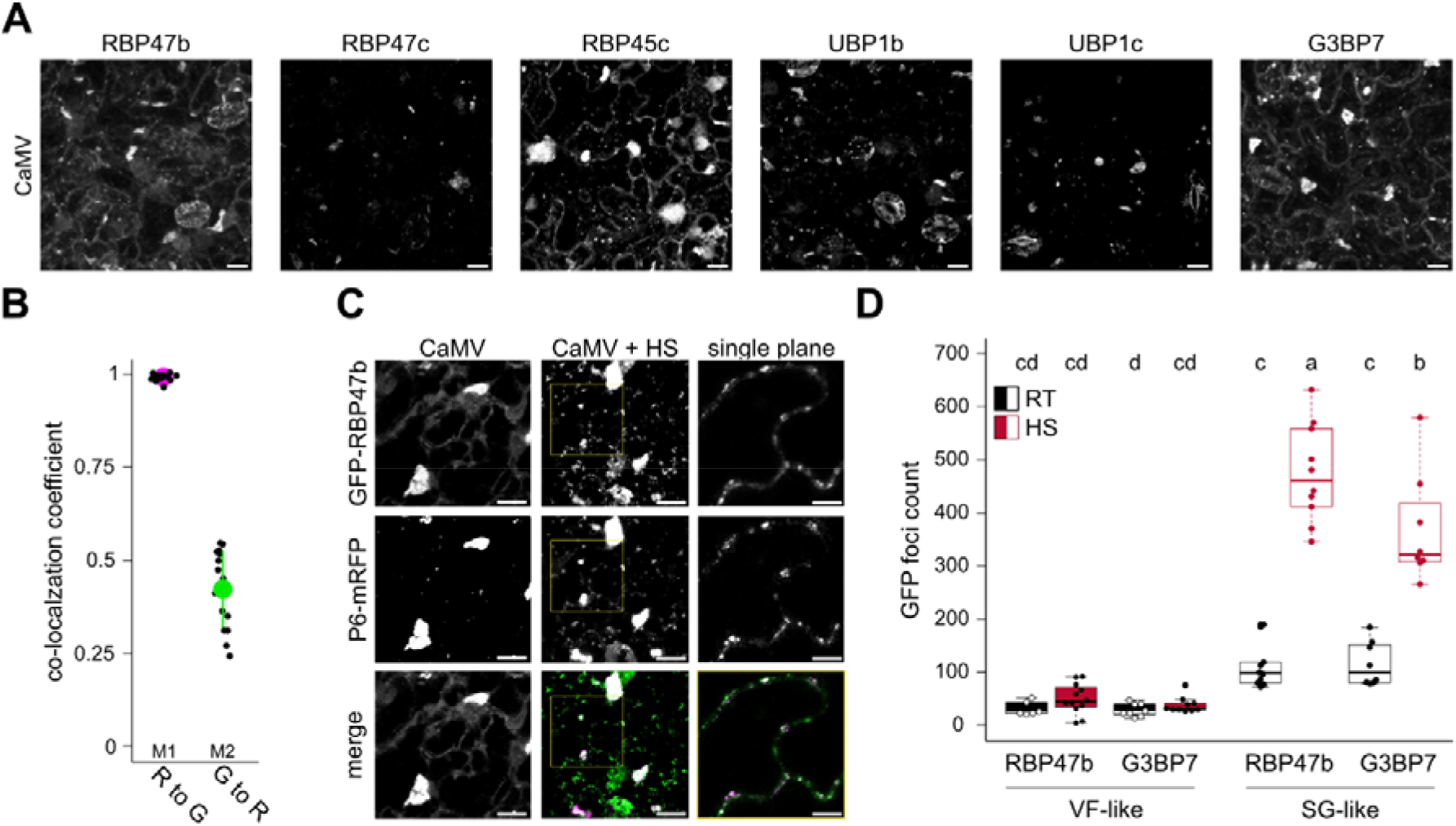
Stress granule proteins localize to viral factories. **(A**) Localization of canonical SG markers 21 dpi with CaMV. Representative images are confocal Z-stack projections (Scale bars = 10 μm). **(B)** Mander’s co-localization coefficients of GFP-RBP47b (G) and P6-mRFP (R) 21 dpi with CaMV. Values were calculated from 16 z-stacks using the ImageJ plugin JACoP. **(C)** Co-localization of GFP-RBP47b and P6-mRFP 21 dpi with CaMV after 30 min of heat shock (HS). Representative images are confocal Z-stack projections (Scale bars = 10 μm). The inset from CaMV + HS is shown to the right and represents single plain images (Scale bars = 5 μm). **(D)** Foci counts in 100 × 100 μm^2^ in infected tissues for GFP-RBP47b and GFP-G3BP7. Foci were separated into SG-like (<2 μm^2^) or VF-like (>2 μm^2^). Counts were averaged from ten replicate images with a custom ImageJ pipeline. Letters indicate statistical groups determined by one-way ANOVA followed by Tuckey’s HSD test (α = 0.05).

We also analyzed the previously described eIF4A-GFP line reported to assemble SGs in response to heat (Hamada et al., 2018). We confirmed that eIF4A-GFP forms numerous foci in the cytoplasm upon heat shock, however it did not localize to VFs upon CaMV infection (supplemental Figure 2B). Intriguingly, when CaMV infected eIF4A-GFP plants were heat stressed, this marker did enter VFs, morphologically clearly different from the heat shock-induced foci in non-infected plants (supplemental Figure 2B). These results expand the repertoire of VF localized RNA granule proteins, yet the specific absence of eIF4A from VFs could indicate selectivity of SG components by VFs, similar as we observed a specific absence of the PB component DCP1 (Hoffmann et al., 2022), or that eIF4A is a conditional SG component. To evaluate this, we chose arsenite treatment as another commonly used stressor leading to stalled translation and SG assembly (Bernstam and Nriagu, 2000; Sorenson and Bailey-Serres, 2014). We detected polysome disassembly in response to both heat and arsenite, albeit milder for arsenite (supplemental Figure 2C). This quantitative difference on polysomes was also reflected by the more numerous induction of SGs by heat than arsenite in the G3BP7 and RBP47b marker lines (supplemental Figure 2D and E). Notably, eIF4A did not form SG foci upon arsenite stress (supplemental Figure 2D and E), suggesting that eIF4A is a conditional SG component and that SG component composition within VFs in ambient temperatures is closer to arsenite-induced SGs, than heat induced SGs.

It is interesting that CaMV induces SG-like foci beside the VFs, despite enhanced translation levels in infected tissues (Figure 2A and D) (Hoffmann et al., 2022), being opposite to the canonical anti-correlation existing between these two processes. Importantly, the number of SG-like foci increased substantially in infected tissues after heat shock, while VF-like foci remained at constant numbers (Figure 2D), suggesting that infected tissue is competent of *de novo* SG assembly. However, despite the resemblance of these newly assembled heat induced SGs to those observed in non-infected tissues, a closer inspection show that they frequently contained also P6 (Figure 2C) and may therefore be functionally diverted and controlled by CaMV. Taken together, we conclude based on recruitment of several SG proteins, that VFs act as a sponge for RNA granule proteins during CaMV infection, although some selectivity and environmental dependence can be observed as judged from eIF4A.

### VFs, unlike PBs and SGs, do not depend on polysomal mRNA supply

A hallmark characteristic of canonical PBs and SGs is their sensitivity to cycloheximide (CHX), a drug that inhibits translational elongation and thereby locks ribosomes on RNA, inhibiting both ribosome runoff and thereby reducing the available mRNA for granulation. Accordingly, we could observe both the disassembly of DCP5-GFP positive PBs, as well as prevention of arsenite-induced GFP-RBP47b SGs in the presence of CHX (Figure 3A-C). In contrast, this treatment had no evident effect on P6-GFP foci number or signal intensity (Figure 3D and E). This suggests that P6 condensates formed in the absence of infection are not similarly dependent on mRNA supply from ribosomes as canonical mRNA granules. We then used the double marker line GFP-RBP47b and P6-mRFP to test the susceptibility of SG proteins within viral factories to CHX. Like P6 condensates, VFs appeared unaffected by CHX treatment (Figure 3F and G). While small cytoplasmic foci readily disappeared in infected tissue after CHX treatment, the large VFs persisted (Figure 3H) and were still marked by GFP-RBP47b with the same signal intensity normalized to P6 as in the EtOH control (Figure 3I). We then used the SG-inducing conditions heat and arsenite in conjunction with fluorescence intensity monitoring of GFP-tagged RBP47b or G3BP7 in VFs but could not detect any apparent differences between the treatments (Figure 3J). We observed the same behavior with the PB marker DCP5, where CHX treatment diminished canonical PBs but not VFs (Figure 3K). Likewise, neither CHX, arsenite or heat affected the amount of DCP5 fluorescence in VFs (Figure 3L and M). Altogether, this would suggest that VFs along with its SG and PB components are not in a similar interdependence with mRNA channeling from the translation machinery as canonical SGs or PBs.

**Figure 3:**
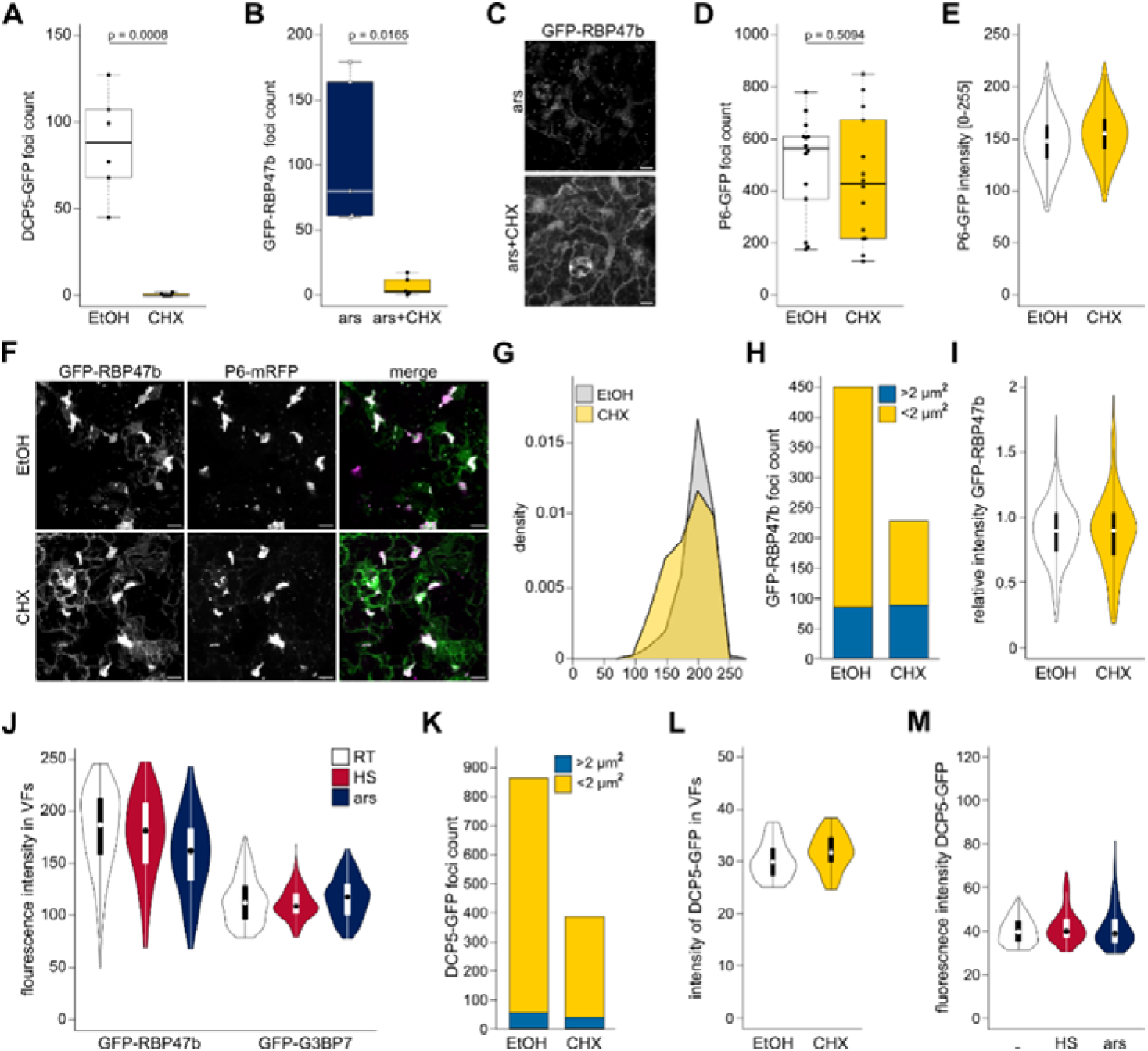
RNA granule components in VFs are unresponsive to SG/PB inhibition or induction. **(A**) DCP5-GFP foci counts in 100 × 100 μm^2^ after EtOH (control) or 200 μM CHX treatment for 2h. Counts were averaged from six replicates. **(B)** GFP-RBP47b foci counts in 100 × 100 μm after 1 mM arsenite + EtOH and 1 mM arsenite + 200 μM CHX treatment for 2h. Counts were averaged from six replicates. **(C)** Representative image of GFP-RBP47b marker line after induction by arsenite (upper panel) and additional treatment with CHX (lower panel) corresponding to (B) (Scale bars = 10 μm). **(D)** P6-GFP foci counts in 100 × 100 μm^2^ after EtOH or 200 μM CHX treatment for 2h. Counts were averaged from 14 replicates. **(E)** Fluorescent intensity of P6-GFP foci after EtOH and CHX treatment in (D). Violin plots represent counts of 7513 (EtOH) and 5839 (CHX) foci. (**F**) Representative image of GFP-RBP47b and P6-mRFP double marker line 21 dpi with CaMV after treatment with either EtOH (upper panel) or 200 μM CHX (lower panel) for 2h (Scale bars = 10 μm). **(G)** Frequency diagram of P6-mRFP signal intensity in viral factories after EtOH or CHX treatment. X-axis denotes the fluorescence intensity, the y-axis the counts in each bin (binwidth = 25). The same imaging set up was used as in (F, G, H, I) **(H)** GFP-RBP47b total foci count split between SG-like foci (<2 μm^2^) and VF-like foci (>2 μm^2^) after EtOH or CHX treatment. The same imaging set up was used as in (F, G, I). **(I)** Relative intensity of GFP-RBP47b compared to P6-mRFP within viral factories after EtOH or CHX treatment. The same imaging set up was used as in (F, G, H). **(J)** Fluorescent intensity of GFP-RBP47b and GFP-G3BP7 foci in ambient temperatures, after 30 min heat shock at 38C° or 1 mM arsenite treatment for 2h. n= 135 – 153 VFs in each condition **(K)** DCP5-GFP total foci count split between SG-like foci (<2 μm^2^) and VF-like foci (>2 μm^2^) after EtOH or 200 μM CHX treatment for 2h. **(L)** Fluorescent intensity of DCP5-GFP in VFs after EtOH (n =57) or CHX treatment (n=38) as in (K). **(M)** Fluorescent intensity of DCP5-GFP in VFs in ambient temperatures, after 30 min heat shock at 38C° or 1 m M arsenite treatment for 2h. n= 82 – 95 VFs in each condition. Statistical significance for A,B and D was calcula ted by Welch Two Sample t-test.

### RNA granule proteins remain highly mobile in viral factories

The insensitivity of SG proteins to CHX regarding their association with VFs could point to another role than translational repression of transcripts. Furthermore, the unconventional overlap of PB and SG components may indicate an aberrant RNA granule character that is usually associated with relative immobility of RNA granule proteins compared to SGs and PBs in liquid phase (Frydryskova et al., 2020). However, FRAP analysis showed that G3BP7, RBP47b and RBP45c were all highly mobile within VFs and recovered within 5-10s after bleaching (Figure 4A and B). Furthermore, these proteins were also constantly exchanged with the surrounding cytoplasm, as indicated by abundant fluorescence recovery after bleaching the whole VF (Figure 4C). Thus, SG proteins are highly dynamic within VFs, and shows comparable FRAP recovery rates to that observed in mammalian SGs for G3BP7 and RBP47b homologues G3BP1 and TIA-1, respectively (Kedersha et al., 2000; Kedersha et al., 2005).

**Figure 4:**
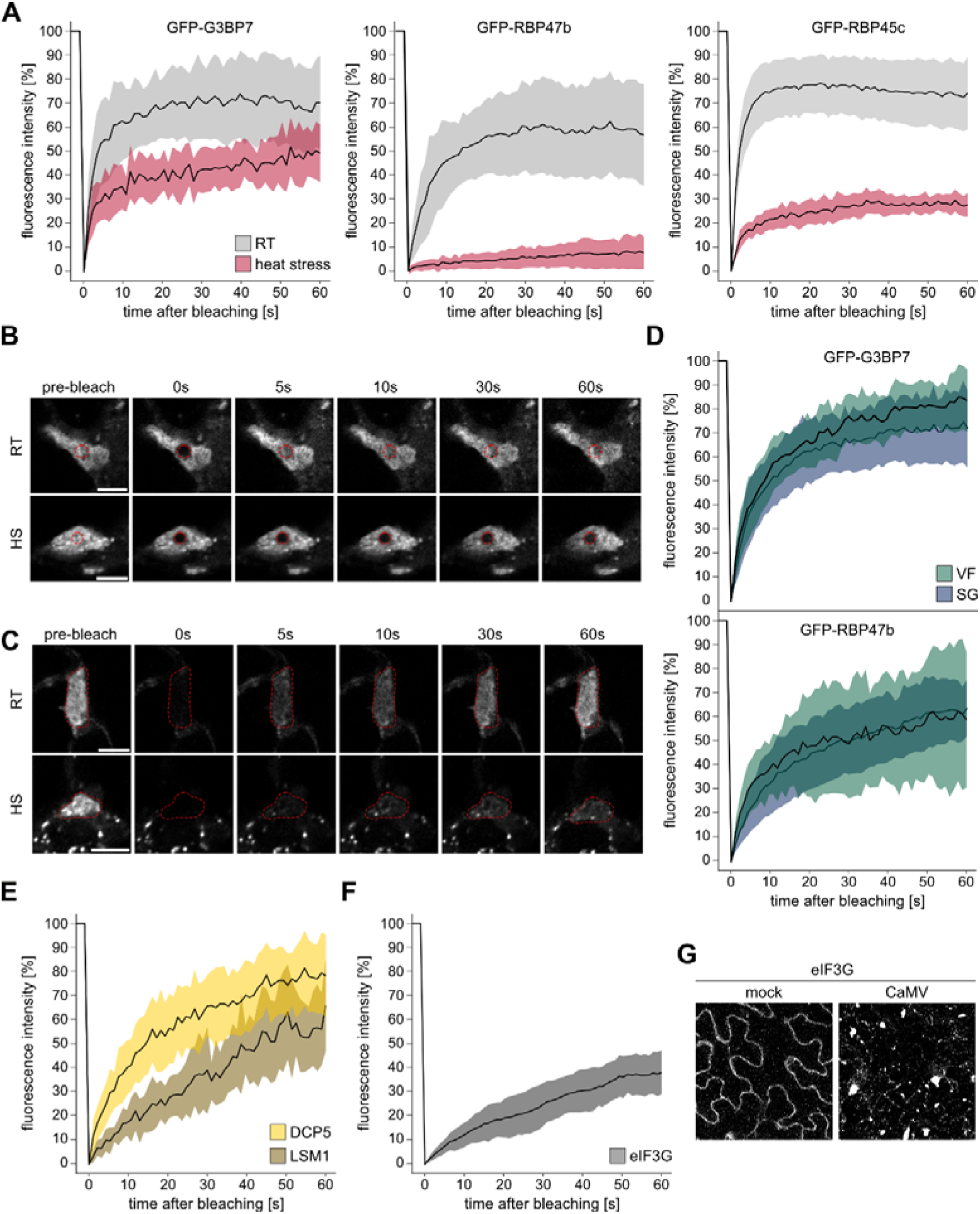
RNA granule proteins shuttle rapidly within VFs and between their surrounding. **(A**) FRAP analysis of indicated proteins in viral factories at 21 dpi at ambient temperatures (RT) or after 30 min of 38 °C heat shock (HS). Normalized fluorescence intensities are plotted against time after photobleaching. n=5-13 **(B)** Representative image series from FRAP analysis of GFP-RBP47b after photobleaching corresponding to RT and HS in (A). Photobleached region is indicated by a red circle. Scale bars = 5 μm. **(C)** Representative image series from FRAP analysis of GFP-G3BP7 after photobleaching the whole viral factory at RT or after 30 min of 38 °C HS. Photobleached region is indicated by a red outline. Scale bars = 5 μm. **(D)** FRAP analysis of indicated proteins in viral factories (VF) and stress granules (SG) at 21 dpi after 2h of 1mM arsenite treatment. Normalized fluorescence intensities are plotted against time after bleaching. n=11/13 for VFs and 17/31 for SGs **(E)** FRAP analysis of DCP5 (n=35) and Lsm1a (n=16) indicated proteins in viral factories at 21 dpi at ambient temperatures. Normalized fluorescence intensities are plotted against time after bleaching. **(F)** FRAP analysis of eIF3g in viral factories at 21 dpi at ambient temperatures. Normalized fluorescence intensities are plotted against time after bleaching. n=7 (**G**) Localization of eIF3g-GFP under uninfected mock conditions and 21 dpi with CaMV. Representative images are composed of confocal Z-stacks (Scale bars = 10 μm). **A, D-F** Solid lines represent mean, shades denote ± standard deviation.

To further estimate analogy between VFs and canonical SGs, we compared their component dynamics during heat and arsenite stress. Upon heat application G3BP7 still recovered quickly in both VFs and SGs, although a fraction of the protein became immobile (Figure 4A; supplemental Figure 4A). In contrast, RBP47b, RBP45c and eIF4A were largely immobile within VFs and SGs in response to heat (Figure 4A and B; supplemental Figure 4). We conclude that SG component dynamics is different in default VFs compared to heat induced SGs, but VFs undergo an intriguing transformation in this direction when heat stressed as already supported by the conditional eIF4A targeting (supplemental Figure 2B).

Considering that eIF4A foci assemble only upon heat but not arsenite stress in contrast to G3BP7 and RBP47b and, that SGs induced by heat and arsenite can differ in many aspects (Frydryskova et al., 2020), we also performed FRAP analysis of these components after arsenite treatment. Notably, both proteins remained mobile and recovered quickly within both VFs and SGs in contrast to their behavior after heat shock (Figure 4D). The two canonical PB proteins LSM1 and DCP5 were also highly mobile within the VF, although LSM1 recovery was slower, and the protein had a larger immobile phase than DCP5 (Figure 4E). The rapid shuttling of the tested RNA granule proteins within the VFs and between VFs and cytoplasm suggests that large fractions of them do not bind strongly to the immobile P6 matrix. We finally used eIF3g known to directly interact with P6 (Park et al., 2001) to evaluate its presence in VFs and more importantly, whether a direct interaction leads to a similar immobility as observed for P6. eIF3g-GFP localized prominently to VFs and indeed displayed slower recovery than especially SG proteins but was still clearly mobile compared to P6 (Figure 4G and H). Altogether, we conclude that i) RNA granule proteins are not rigidly bound to or aggregated within the immobile P6 phase, but shuttle between the VFs and their surroundings ii), VF component composition and mobility resembles arsenite but not heat SGs and iii), that VF components adapt heat SG characteristic dynamics upon exposure to this stress.

### The *35S* genomic viral RNA binds PB while avoiding SG components *in planta*

An outstanding question was whether SG components participate in CaMV infection as previously established for PB components DCP5 and Lsm1a/b (Hoffmann et al., 2022). The 9 members of the RBP47b gene family are largely uncharacterized (for phylogenetic tree see (Sorenson and Bailey-Serres, 2014)), with knock-out phenotypes identified individually for *UBP1b* and *UBP1c* suggesting some non-redundant functions (Bailey-Serres et al., 2009; McCue et al., 2012). We established a collection of T-DNA insertion mutants in members of this gene family as well as a triple mutant *(rbp47a ubplb ubplc)* to address their importance for CaMV accumulation, but they had no evident effect as even the slight reduction initially observed in *ubplb* and *ubplc* was absent in the combinatorial triple mutant (Figure 5A). Possibly, there is a high degree of redundancy within this gene family and their functions in CaMV infection, as several of them localize to VFs (Figure 2A), or they are not influencing virus accumulation despite their VF localization.

**Figure 5:**
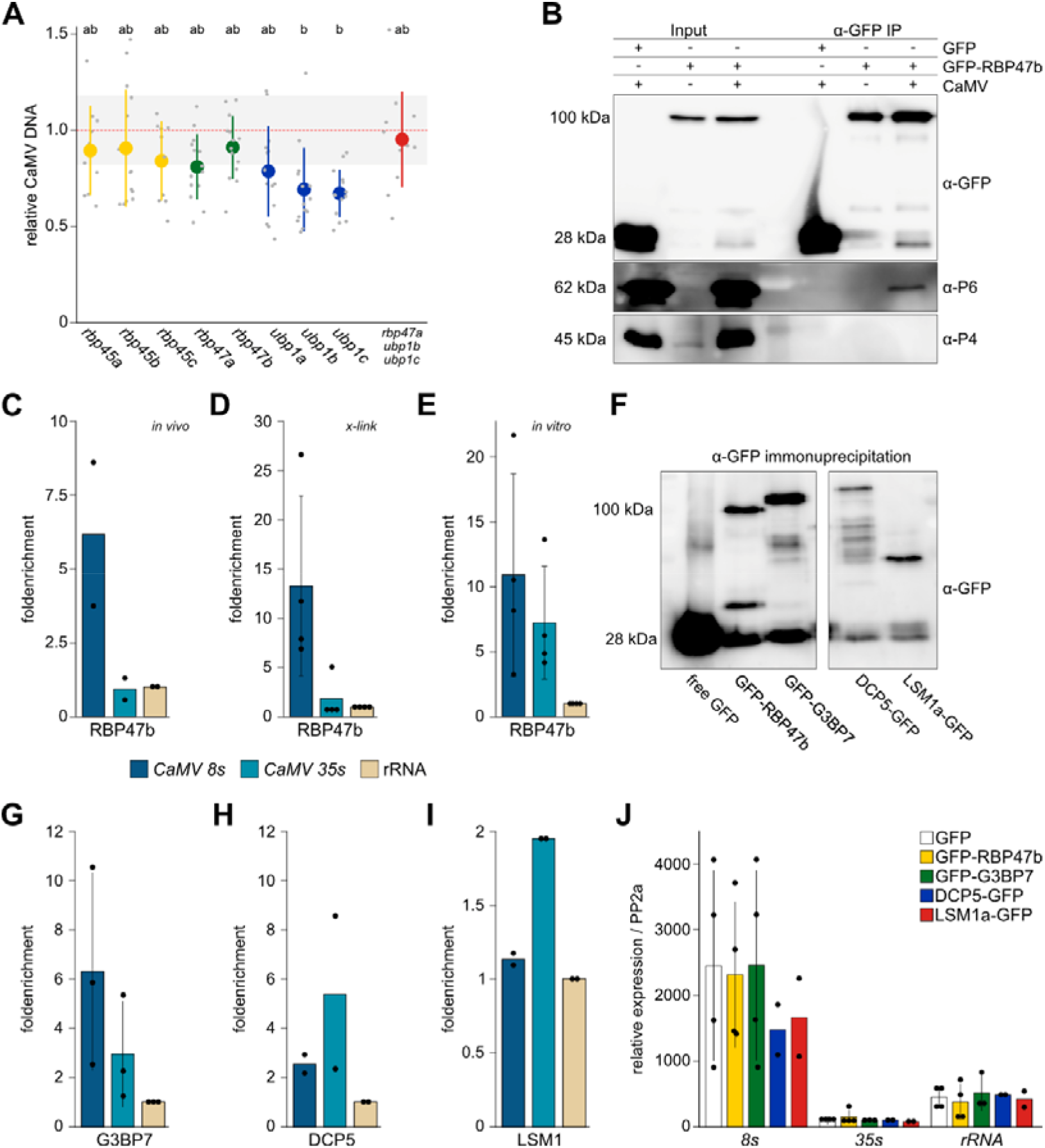
SG and PB proteins gain different access to viral RNA. **(A)** Viral DNA accumulation in systemic leaves of indicated genotypes at 21 dpi, determined by qRT-PCR. Average is depicted by one large, colored dot and replicates by small grey dots (n = 12) relative to Col-0 plants and normalized to *18S* ribosomal DNA as the internal reference. Letters indicate statistical groups determined by one-way ANOVA followed by Tuckey’s HSD test (α = 0.05). **(B**) Co-immunoprecipitation analysis of viral proteins P6 and P4 with GFP-RBP47b at 21 dpi with CaMV. Infected plants expressing free GFP and non-infected GFP-RBP47b plants functioned as controls. **(C)** Fold enrichment of viral RNAs in native RNA-immunoprecipitation assay (RIPA) from GFP-RBP47b over free GFP expressing plants using *ribosomal (r)RNA* for calibration. Data points represent independent experiments. **(D)** As (C) but including an *in planta* formaldehyde cross-linking step prior to RIPA. **(E)** Fold enrichment of viral RNAs in an *in vitro* RIPA with GST-RBP47b over the GST control using *rRNA* for calibration. Data points represent independent experiments. (**F)** Western blot analysis using anti-GFP to verify capture of baits in the RIPAs of GFP, RBP47b, G3BP7, DCP5 and Lsm1a. **(G, H** and **I)** Fold of enrichment of viral RNAs in formaldehyde cross-linked RIPAs from GFP-G3BP7 (G), DCP5-GFP (H) and Lsm1a-GFP (I) over free GFP expressing plants using *rRNA* for calibration. Data points represent independent experiments. **(J)** Relative expression of viral RNAs and *rRNA* in input fractions of RIPA samples normalized to housekeeping gene *PP2a.*

A primary function of SG and PB components is RNA regulation through RNA binding, prompting us to address if any of these components could be detected in association with viral RNA during infection. First, we infected GFP-RBP47b and free GFP control plants and harvested symptomatic tissue at 21 dpi for GFP-based co-precipitation. We detected co-purification of viral protein P6 but not P4 using western blot analysis (Figure 5B), and interestingly also the viral *8S* leader RNA but not the protein-coding full viral *35S* RNA or control *rRNA* (Figure 5C). To reduce the risk of disassociation during isolation, we included an *in planta* formaldehyde cross-linking step which led to slightly better capture of *8S* but still no detectable enrichment of *35S* (Figure 5D). An *in vitro* association assay developed largely according to (Dember et al., 1996) showed, however, that GST-RBP47b could bind *35S* RNA with comparable efficiency to *8S*, both being highly enriched over the GST control and in comparison to our background *rRNA* control (Figure 5E). Next, we extended the *in planta* RNA immunoprecipitation assay (RIPA) to additionally include G3BP7, DCP5 and Lsm1a. This revealed that G3BP7 was also associated with *8S* above *35S* RNA similar to RBP47b, while both PB components oppositely enriched *35S* above *8S* (Figure 5F-H). A western blot verified capture of the baits in RIPA (Figure 5I) and total transcript levels in the assay (Figure 5J) suggested that while SG components may associate with *8S/35S* in a somewhat total quantity-dependent manner, PB components DCP5 and Lsm1 select for and thus appear as more specific regulators of *35S* RNA.

### P6 can stabilize polysomes as well as localize to and suppress SGs via eIF3g

As observed in Figure 2C, P6 localized to potentially newly induced SGs upon heat stress in infected tissue, but whether these were truly *de novo* assemblies and in particular if this also occurred in the absence of infection remained unknown. Marker-lines for GFP-RBP47b and GFP-G3BP7 expressing P6-mRFP were challenged with arsenite or heat followed by quantification of SG-like foci assembly. Intriguingly, the P6-mRFP lines generally showed strongly decreased amounts of detectable Rbp47b and G3BP7 foci compared to their parental lines (Figure 6A), except for CMI line #1 during heat. The signal of GFP-RBP47b appeared overall low in the P6 lines, but a western blot could not detect any clear difference here or in GFP-G3BP7 levels (supplementary Figure 4). Notably, those residual SG foci that still formed, were frequently co-labelled by P6 (Figure 6B), supporting that P6 can localize to SGs also in the absence of infection.

**Figure 6:**
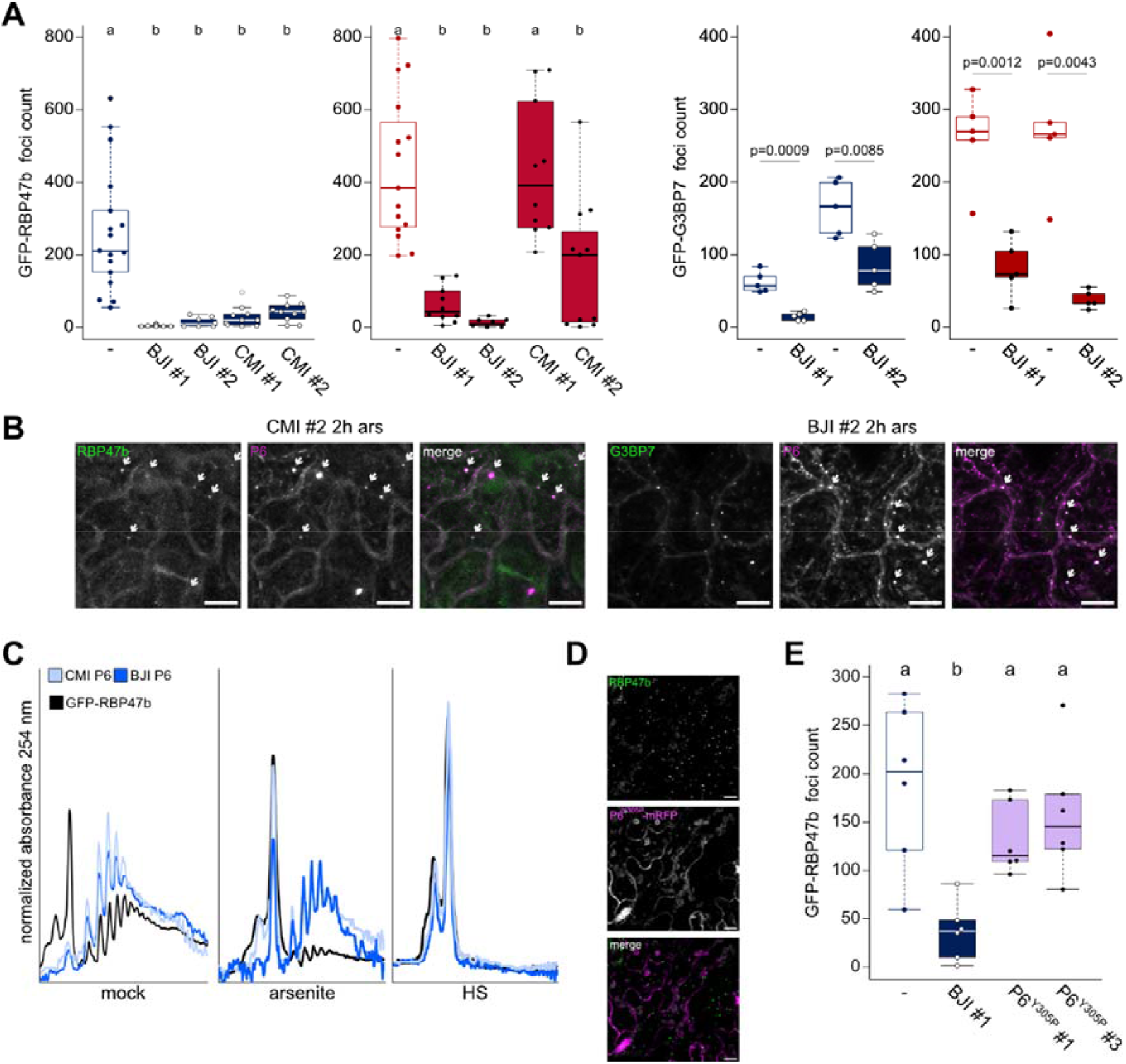
P6 inhibits Stress Granule formation. **(A**) GFP-RBP47b and GFP-G3BP7 foci counts in double marker lines with P6-mRFP in 100 × 100 μm^2^. Counts were averaged from 10 – 15 replicates after 1 mM arsenite treatment for 2h (left panel, blue) or 30 min of 38 °C heat shock (HS) (right panel, red). # denotes independent transgenic lines. The G3BP7 data with line #1 and #2 were done separately and thus have individual controls (-) to their left, statistical significance for these groups were determined by Welch Two Sample t-test. **(B)** Representative image of GFP-RBP47b and GFP-G3BP7 corresponding to A and B. Examples of condensates containing both the SG marker and P6 are marked by white arrows. (Scale bars = 10 μm). **(C)** Polysome profiles of GFP-RBP47b with and without expression of P6s in untreated and after arsenite or heat treatment as in (A). **(D)** Representative image of GFP-RBP47b and P6Y305P-mRFP double-marker line (Scale bars = 10 μm). Note, the signal intensity of this mutant P6 was clearly lower, causing a partial bleed-through signal from chloroplasts in the P6 magenta channel. (**E**) GFP-RBP47b foci counts in double marker lines with P6Y305P-mRFP in 100 × 100 μm^2^. Counts were averaged from six replicates after arsenite treatment. Two independent lines were used (#1 and #2). **(A,E)** Letters indicate statistical groups determined by one-way ANOVA followed by Tuckey’s HSD test (α = 0.05).

CaMV P6 is a master regulator of *35S* RNA translation (Pooggin and Ryabova, 2018), including mechanisms of translation re-initiation together with eIF3g (Park et al., 2001). Excitingly, we found that the polysome to monosome ratios were clearly increased in P6-mRFP lines compared to the parent (Figure 6C), suggesting that P6 may have a global impact on polysomes and could be a contributor to the observed increase in polysomes during CaMV infection (Park et al., 2001; Hoffmann et al., 2022). Moreover, arsenite strongly reduced polysomes in the parental control, while P6-mRFP lines largely resisted this despite an evident increase in monosomes and, heat stress was sufficient to fully disassemble polysomes in all lines (Figure 6C). While it is conceivable that polysome stabilization by P6 contributes to SG inhibition upon arsenite treatment, the strong inhibition of SGs despite full polysome disassembly during heat stress would point towards an additional uncoupled mechanism. We pursued the importance of P6 translation re-initiation mechanisms in SG inhibition by establishing GFP-RBP47b lines expressing P6-mRFP with tyrosine 305 (P6Y305P) swapped to proline, a mutation that disrupts the essential eIF3g interaction and translation transactivation *in vitro* (Park et al., 2001) but retains e.g. suppression of salicylic acid responses (Love et al., 2012). The Y305P mutation compromised P6 suppression of SGs in response to arsenite, removed co-localization and also failed to show any self-condensates (Figure 6D and E). Together, this established the capacity of P6 to counteract SG amounts during stress, its prominent localization to SGs and the potential importance of eIF3g interaction in these processes.

### SG inhibition and trans-activation can be uncoupled and are reduced by P6 condensation

Some animal viruses sequester SG components to dampen their response (Emara and Brinton, 2007; Panas et al., 2012) and analogously, targeting of SG components to VFs could in a similar scenario negatively impact SGs. However, because the fluorescence intensity of GFP-RBP47b was much lower in the P6-mRFP mock condensates than in VFs (Figure 7A), this appeared opposite to SG inhibition (Figure 6A vs. Figure 2D). A particular feature of P6-mRFP is reduced self-condensation and increased solubility compared to a P6-tagRFP fusion that showed much larger self-condensates and no soluble signal (Figure 7B). Using fractionation by differential centrifugation, their difference in solubility was further supported (Figure 7C) and P6-mRFP soluble to condensate ratio was close to that observed in infected tissue (Figure 7D). We utilized this solubility difference to address how P6 condensation and associated SG component sequestration contributes to SG inhibition. P6-mRFP and P6-tagRFP were co-expressed with GFP-RBP47b in *Nicotiana benthamiana* and heat-induced SGs were quantified. While both P6s reduced the amount of SGs, this effect always appeared stronger for the more soluble P6-mRFP (Figure 7E).

**Figure 7:**
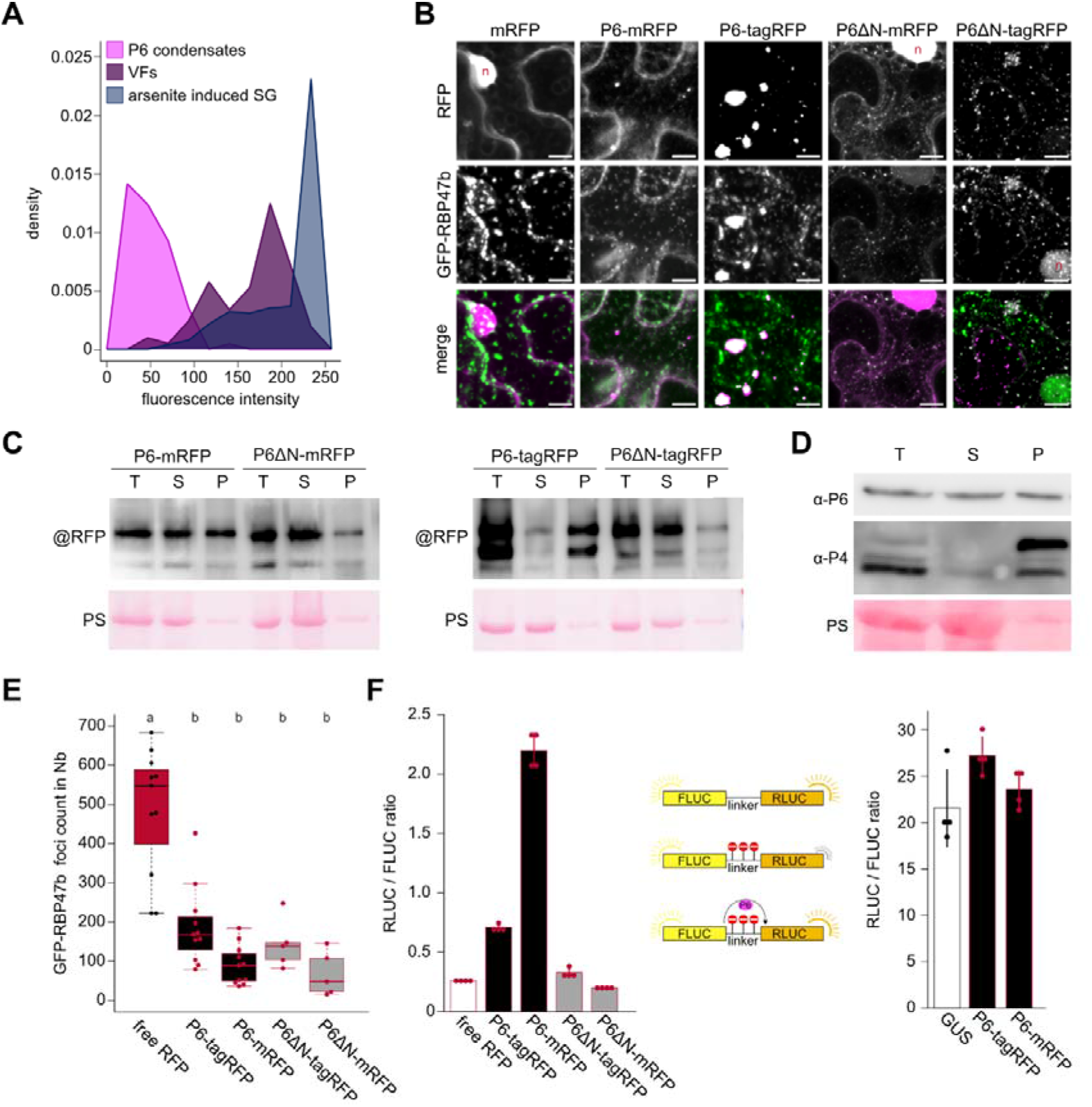
Soluble P6 can inhibit SGs regardless of capacity to trans-activate translation. **(A)** Density polygon (binwidth = 25) of GFP-RBP47b fluorescence intensities within pre-assembled P6-mRFP condensates (n=90) in the transgenic lines (Figure 6A), viral factories (VFs) from infection at 21 dpi (n=83) and SGs induced using 1 mM arsenite treatment (n=634). **(B)** Representative image composed of confocal z-stack projection of GFP-RBP47b / P6 co-expressions in *N. benthamiana* at 3 days after infiltration. The specific P6 constructs are indicated above the images. (Scale bars = 10 μm). **(C)** Western blot analysis of the P6 constructs expressed in *N. benthamiana* at 3 dai. Total (T) protein samples were extracted and subjected to differential centrifugation resulting in soluble (S) and pellet (P) fraction. Blots were probed with mRFP and tagRFP specific antibodies, respectively. Ponceau S (PS) staining served as loading control. **(D)** Western blot of CaMV proteins P6 and P4 in systemically infected Arabidopsis leaves at 21 dpi probed with specific antibodies. Total (T) protein samples were extracted and subjected to differential centrifugation resulting in soluble (S) and pellet (P) fraction. PS staining served as loading control. **(E)** GFP-RBP47b foci counts in 100 × 100 μm^2^ of *N. benthamiana* leaves co-expressing indicated P6 constructs at 3 dai after a 30 min 38 °C heat shock. Counts were averaged from ten (full length constructs) or five (P6ΔN constructs) replicates and analysed with a custom ImageJ pipeline. **(F)** Analysis of transactivation activity for indicated P6 constructs compared to free mRFP in *N. benthamiana*. P6s and control were co-infiltrated with FLUC-3Xstop-RLUC and luciferase activity was analyzed at 3 dai (left panel). Increased activity of transactivation is indicated by a higher RLUC to FLUC ratio (n=4). P6 has no influence on the ratio of RLUC/FLUC when co-expressed with FLUC-linker-RLUC without the stops (right panel; n=4).

To further strengthen that the soluble P6 pool is stronger in suppressing SGs, we explored a 17 amino acid deletion in the N-terminus region known to be essential for P6 condensation (P6Ndel3-20, hereafter referred to as P6ΔN) (Haas et al., 2008; Laird et al., 2013), and fused this P6 mutant to both tagRFP and mRFP. As expected, the solubility clearly increased for both fusions (Figure 7B and C). The P6ΔN mutant retained the capacity to suppress heat SGs when co-expressed with GFP-RBP47b in *N. benthamiana* and resided at the level of the WT protein (Figure 7E). As we initially hypothesized that P6 may suppress SGs by both translation dependent and independent mechanisms, we assessed the essential activity of P6 and mutants to transactivate translation of consecutive open reading frames separated by stop codons in the translation reporter FLUC-3xSTOP-RLUC construct (Figure 7F). P6 co-expression does not alter FLUC to RLUC ratios when expressed with the control FLUC-linker-RLUC plasmid (Figure 7F). We found that P6-mRFP was a much stronger translational transactivator than P6-tagRFP and, that the P6ΔN mutants were incapable of transactivation regardless of the RFP tag. These results revealed a previously undescribed importance for the N-terminus of P6 for *in planta* transactivation. Even more importantly, P6ΔN retained the capacity to suppress SGs but not transactivation. This at least partially uncouples these mechanisms to support a translation-independent function of P6 in SG suppression, which we anticipated from the suppression observed also in response to heat with full polysome disassembly (Figure 6). However, these results do not rule out a translation dependent mechanism that very likely contributes SG suppression by stability of polysomes during arsenic stress in P6-mRFP transgenic lines (Figure 6). Because condensation was reduced but not SG suppression for P6ΔN, we propose that soluble P6 promotes SG suppression and P6 condensation is likely to reduce this function as seen in P6-tagRFP.

## DISCUSSION

Membrane-free condensation of biomolecules into larger entities is a ubiquitous event including many distinct higher-order complexes of RNA and regulative proteins. Despite recognition of the phenomenon for a long time, the purpose of the actual condensation process has remained largely hypothetical. In the current work, we focused on the specific condensate of CaMV viral factories (VFs) orchestrated by the multifunctional P6 protein. Early work established that VFs lack enclosing membranes and are mainly composed of RNA and protein with viral particles dispersed within (Martelli and Castellano, 1971). For long, VFs were believed to not exchange much content with the surrounding (Kitajima et al., 1969; Conti et al., 1972), a view that was overthrown by the intriguing finding that viral particles are mobilized from within VFs during conditions mimicking aphid infestation (Bak et al., 2013). Considering that CaMV particles are 50 nm in diameter, the VFs should be a highly flexible matrix for them to mobilize, despite our finding that P6 itself is largely static. SG and PB proteins on the other hand, showed very fast mobilities both within and between VFs and the surrounding. This supports that VFs are dynamic structures, with G3BP7 and RBP47b showing comparable mobility to their mammalian homologues in arsenite-induced liquid-liquid phase-separated (LLPS) SGs (Buchan and Parker, 2009). Intriguingly, these dynamics change when VFs are subjected to heat but not arsenic stress, including prominent recruitment of the heat-specific SG component eIF4A. The composition of LLPS organelles can change rapidly in response to stress (Moore et al., 2011) as we observed for eIF4A in VFs. VFs grow in size and reduce in numbers over time, suggesting growth by fusion, a common behavior of phase separated condensates (Alberti et al., 2019). Furthermore, P6-GFP condensates formed by transient expression in *N. benthamiana* were recently reported to partially respond to the LLPS disruptive agent 1,6-hexanediol (Alers-Velazquez et al., 2021). Interestingly, there seems to be a wide range of variation among LLPS structure morphologies and properties (Fare et al., 2021) and, e.g. mammalian SGs are frequently irregular in shape with substructures (Souquere et al., 2009). The amorphous shape and presence of lacunae suggests that VFs also have substructures that furthermore varies to some extent between hosts and viral strains (Schoelz and Leisner, 2017; Hoffmann et al., 2022). As we find that VFs contain several SG and PB components, they appear as more general melting pots for RNA metabolic proteins and diverted from these otherwise canonically distinct LLPS condensates. The composition of LLPS compartments could be largely controlled by central scaffolding protein nodes forming multivalent interaction networks (Fare et al., 2021). Notably, P6 is a truly multivalent node regarding several LLPS criteria; i) contains three described RNA binding domains ii), shows complex self-association involving at least four distinct domains and iii), binds directly to a multitude of proteins including eIF3g and VCS that we found in VFs (Park et al., 2001; Schoelz and Leisner, 2017; Lukhovitskaya and Ryabova, 2019; Hoffmann et al., 2022). Scaffolding protein nodes would have reduced mobility compared to recruited clients owing to their multiple interactions in the network (Fare et al., 2021), as we observed here for P6 in relation to all other assessed VF components. Analogously, TSN2 serving as a docking scaffold for SG assembly in plants also display very slow mobility in the condensates compared to e.g., RBP47b (Gutierrez-Beltran et al., 2015; Gutierrez-Beltran et al., 2021). As part of its essential role in CaMV translation, P6 interactions include ribosomal proteins L13, L18 and L24 (Leh et al., 2000; Park et al., 2001; Bureau et al., 2004) and regulators eIF3g (Park et al., 2001) and TOR (Schepetilnikov et al., 2011). Taking VF coating with ribosomes (Shepherd, 1976) together with our finding of eIF3g being extensively localized to VFs, we speculate that VFs may serve a function as reservoirs of CaMV translation supporting factors. We conclude that many observations support the possibility that LLPS participates in VF condensates, and overall, they seem to provide a highly dynamic environment for viral and host factors.

RNA granule proteins were not responsive to CHX treatment within VFs, suggesting that their localization here does not depend on polysomal release of mRNAs similarly as canonical granules (Teixeira et al., 2005; Weber et al., 2008). We found that SG components bound the extremely abundant non-translated *8S* viral RNA. The proposed function for *8S* RNA is to decoy the plant RNA silencing defense through the massive generation of small interfering RNAs targeting the *8S*, but not the genomic *35S* RNA needed for CaMV replication (Blevins et al., 2011; Hohn, 2015). Because *8S* and *35S* RNAs showed comparable binding to RBP47b *in vitro*, we believe that it is rather the non-translatability and shear abundance than sequence specificity that drives the *8S* association with SG components similar to non-translating RNA storage in mammalian cells (Khong et al., 2017). Analogous to decoying the silencing machinery, we speculate that *8S* binding may sequester SG components from their usual RNA clients and thereby suppress their canonical functions. PB components equally localize to the VF matrix, however, they have access to the genomic viral *35S* RNA, plausibly linking them to supporting CaMV translation (Hoffmann et al., 2022).

In multiple instances, animal viruses are able to counteract SG assembly (Lloyd, 2016; Poblete-Duran et al., 2016), and some plant viral proteins have also been found to interact and possibly interfere with core SG components (Krapp et al., 2017; Makinen et al., 2017; Reuper et al., 2021). We found suppressed assembly of SGs monitored via G3BP7 and RBP47b during heat and arsenic stress in plants with ectopic expression of P6 and, that the residual SGs forming frequently contained P6. We consider two nonexclusive modes of SG inhibition by P6: translation dependent, translation independent. Translation dependent involves a global stabilization of polysomes to negatively impact SG numbers, as they depend on non-translating RNAs. In support, the P6Y305P mutant that is incapable of eIF3g interaction and translation transactivation does not suppress or localize to SGs anymore but should be carefully considered owing to its comparably low accumulation levels. Translation independence is suggested as most transgenic P6 lines still show strong SG suppression despite a complete polysome disassembly during heat stress, as well as the P6ΔN mutant capable of SG suppression but not transactivation of translation. The mechanisms by which viral proteins suppress SGs in mammals are diverse (Lloyd, 2016; Poblete-Duran et al., 2016), but the functional analogy between P6 and virus-induced human Adenosine deaminase acting on RNA 1 (ADAR1) is striking. ADAR1 is believed to suppress SGs both translation-dependently and -independently, it localizes to SGs and the double-stranded RNA binding domain is essential for translation-independent inhibition of SGs (Corbet et al., 2021). The double-stranded RNA domain of P6 is known to bind and activate TOR kinase for translation transactivation and autophagy inhibition (Schepetilnikov et al., 2011; Zvereva et al., 2016). Interestingly, TOR signaling can promote SGs in animal models (Sfakianos et al., 2018), outlining it as a potential hub to influence SGs both translation-dependently and - independently upon P6 manipulation. Further dissection of multifunctional P6 is a true challenge that may provide valuable insights into underlying mechanisms of SG inhibition in plants.

Despite strong SG inhibition by ectopic P6, CaMV infected tissue still assembles abundant SG-like foci in response to heat and arsenic stress. Evidently their true SG nature and relatedness to VFs remain unclear owing to a frequent P6 presence but, may suggest differential P6 regulation during infection. By using different protein fusions that influence the solubility of P6, we found that the more soluble P6 version was stronger in both transactivation and SG suppression. As there should be a gradual decrease in the amount of soluble P6 via VF condensation along maturation of infection in a cell, we propose that SG suppression and transactivation may reduce and eventually inactivate through condensation. As P6 is a main symptom determinant, its condensation may also be important according to the idea of self-attenuation (Schoelz and Leisner, 2017), where some plant viruses have been found to limit long-term damage by inactivating their virulence factors (Torres-Barcelo et al., 2008; Zhang et al., 2017; Shukla et al., 2021). For endogenous plant genes, such a dependency was found for ARF condensation in reducing auxin responsiveness (Powers et al., 2019). We conclude this work with a hypothetical working model (Figure 8).

**Figure 8:**
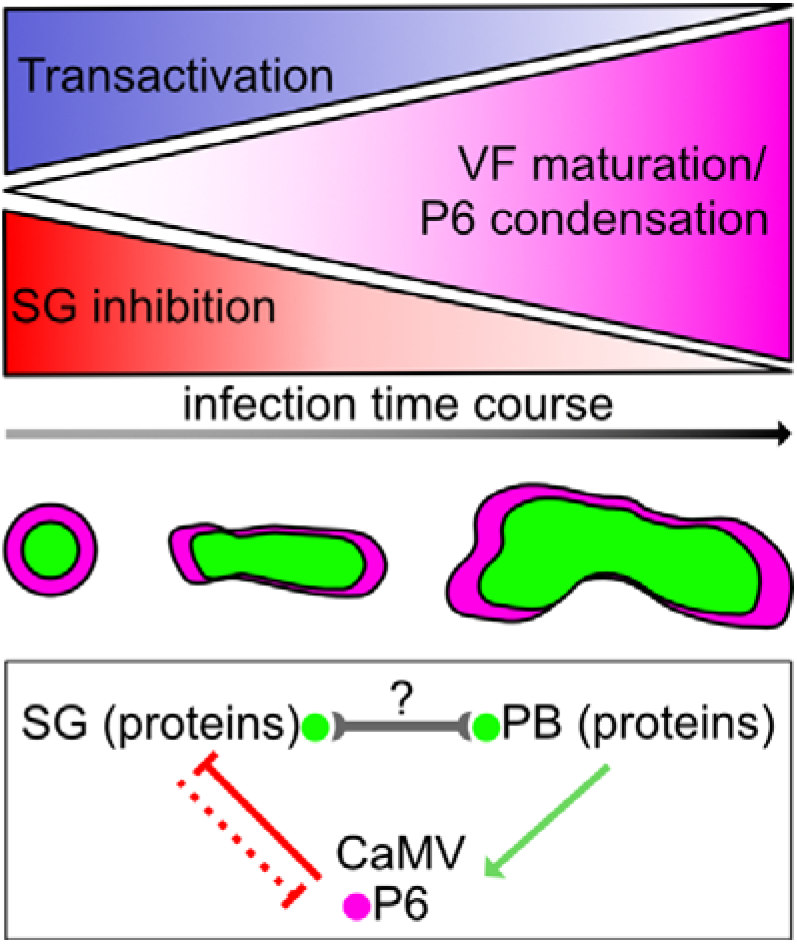
Proposed model of SG-related P6 functions during CaMV infection. P6 condensation within VFs increases during the infection time course, leading to a reduction of P6 functions in transactivation and inhibition of stress granules. SG and PB proteins accumulate inside VFs, where PB components aid the virus translation, while SG components are probably inactivated from their antiviral capacities.

## MATERIAL and METHODS

### Plant Material, Growth Conditions

Arabidopsis and *N. benthamiana* plants were grown in walk-in chambers in standard long day conditions (16h light / 8h dark cycle) at 22°C and 65% relative humidity for crossing, propagation, and transient expression assays. For infection experiments, plants were grown in short day conditions (120 mE, 10h light / 14h dark cycle) at 22°C and 65% relative humidity. All T-DNA and marker lines used in this study were in the Arabidopsis accession Columbia (Col-0) which was taken as control for all experiments.

### Plasmid Construction, Generation of Transgenic Lines and Transient Expression

Clones containing UBP1b, UBP1c, RBP47b, RBP47c, RBP45c, G3BP7 and eIF3g were PCR amplified from Arabidopsis Col-0 cDNA and inserted into the pENTR/D-TOPO cloning plasmid. Entry clones of P6 coding sequences were described before (Hafren et al., 2017) and used for site-directed mutagenesis to obtain Y305P and P6ΔN mutants. The pENTRY clones were recombined into pGWB vectors for mRFP and tagRFP fusions (Nakagawa et al., 2007) and pUBN/pUBC-DEST for GFP fusions (Grefen et al., 2010). Transgenic lines were generated by floral dipping (Clough and Bent, 1998) and, all lines and constructs are listed in supplemental Dataset1 Table S1. The FLUC-RLUC fusions were generated by construction of a pENTRY/D-TOPO clone containing the coding sequence of RLUC preceded by a linker with and without three stop codons, that was further recombined with pMCD32::FLUC (Ustun et al., 2018) to create FLUC-linker-RLUC and FLUC-3Xstop-RLUC reporters for the transactivation assay. For transient expression, *N. benthamiana* leaves were infiltrated with resuspended Agrobacteria (OD 0.2, 10 mM MgCl2, 10 mM MES pH 5.6, 150 μM Acetosyringone) and the constructs analyzed after 72h.

### Virus Inoculation and Quantification

CaMV infections and virus quantification were performed as described (Hoffmann et al., 2022). Viral DNA levels were determined by qRT-PCR and normalized to *18S* ribosomal DNA.

### Ribosomal Profiling

Transgenic GFP-RBP47b with and without additional expression of P6-mRFP fusions from strains CM1841 (CMI) and Cabb-BJI (BJI) were vacuum-infiltrated with 1mM sodium arsenite and kept for 2h, heat shocked for 30 min at 38°C or, left untreated as a control for the treatments, followed by harvesting in lqN and storage at −80°C until processing. Polysome extraction and profile analysis was performed exactly as described before (Hoffmann et al., 2022).

### Pull-down assays

Pull-down of GFP-fusion proteins from plants was carried out by grinding 1 g of leaf tissue in 2 ml of buffer (100 mM Tris pH8, 150 mM NaCl, 20 mM KCl, 2 mM MgCl2, 1% TX-100, 40U Ribolock ml^-2^ and protease inhibitor cocktail, Roche) followed by clearing at 17 000 × g for 10 min at +4C°. For formaldehyde (FA) cross-linking, tissue was vacuum infiltrated with 1% formaldehyde in PBS, cross-linked for 10 min, quenched with 0.125 M glycine PBS and tissue processed as above. Lysates were rotated with 25 μl anti-GFP magnetic agarose beads (Chromotech) for 60 min +4C°, washed 5 times with 1 ml buffer and eluted with Laemmli sample buffer for western blot analysis and Trizol for RNA isolation. For FA cross-linked RNA, input samples and beads were incubated with protease K for 30 min at +50 C° before isolating RNA using Trizol. Isolated RNA was DNAse treated, re-purified and used for cDNA synthesis followed by qRT-PCR. For *in vitro* pull-downs, we generated an RBP47b deletion mutant lacking 100 amino acids from the N-terminus that constitutes the prion domain (Weber et al., 2008) to increase solubility. GST tagged proteins were bound to glutathione sepharose, washed with IP buffer (100 mM Tris pH 7.5, 150 mM NaCl, 20 mM KCl, 1 mM MgCl_2_, 1 mM EGTA, 0.05% IpegalC630 and 40 U Ribolock ml^-2^) and incubated with total RNA extracted from CaMV infected plants diluted in IP buffer rotating for 30 min largely as in (Dember et al., 1996). After washing 6 times with 1 ml of IP buffer, GST-proteins were eluted with 2 mM reduced glutathione followed by Trizol RNA extraction, cDNA synthesis and qRT-PCR.

### Transactivation assay

The FLUC-linker-RLUC and FLUC-3Xstop-RLUC were co-expressed with indicated P6s and controls in *N. benthamiana* by agroinfiltration and harvested 3 dai for analysis. Analysis was performed using the Dual-Luciferase Assay (Promega) according to manufacturer’s instructions.

### Protein detection and fractionation

To detect proteins, we used western blot analysis essentially as described (Hoffmann et al., 2022), with antibodies against GFP (Santa Cruz Biotechnology; sc-9996), tagRFP (Agrisera [RF5R], mRFP (ChromoTek [6G6], P4 (Champagne et al., 2004), and P6 (Schoelz et al., 1991). Secondary antibodies were horseradish peroxidase-conjugated (GE Healthcare; NA934 and NA931).

For fractionation of P6 between soluble and condensates, we homogenized leaf tissue in buffer (100 mM Tris pH8, 150 mM NaCl and 1% TX-100) to obtain the total sample, followed by centrifugation at 17 000 × g for 10 min +4C° to obtain a supernatant with the soluble P6 and a pellet containing the condensate P6. The pellet was reconstituted in the same buffer using a volume corresponding to the original input and, all samples were supplemented with Laemmli sample buffer and used in Western blot analysis.

### IMAGE J

Images were processed with ZEN black software (Zeiss) and ImageJ version 1.53s. For quantification, Z-stacks were Brightness increased and a “Gaussian Blur” filter (sigma=1) applied. A mask was generated through thresholding and foci were analyzed using the “Analyze Particles” tool with the settings (size =0.1-2.0, circularity=0.5-1.0 for stress granules and size= 2.0-inf for viral factories). Colocalization analysis for RBP47b with P6 was performed using the Plugin JACoP (Bolte and Cordelieres, 2006; Bolte and Cordelières, 2006). FRAP analyses were manually processed in ImageJ. Signal intensities in bleached areas were normalized to an unbleached control area at each timepoint to account for general photobleaching. Afterwards time course series were min-max normalized to evade differences in bleaching efficiency and plotted as percent of initial signal vs time.

### DATA ANALYSIS AND STATISTICAL METHODS

Boxplots / violin plots were made with R v4.0.02. The box represents the interquartile range (IQR), the solid lines represent the median. Whiskers extend to a maximum of 1.5 × IQR beyond the box. Data was tested for normality using the Shapiro-Wilk test. Statistical comparisons of two groups were performed by Welch Two Sample t-test with R v4.0.02. One-way ANOVA followed by a post-hoc Tukey HSD test (α = 0.05) was performed with R v4.0.02 and the R-package “agricolae” (Version 1.3-3; https://cran.rproject.org/web/packages/agricolae/index.html). Test statistics are shown in Supplemental Data Set S5.

## ACCESSION NUMBERS

Sequence data from this article can be found in the EMBL/GenBank data libraries under the following accession numbers: LSM1a(AT1G19120), DCP5 (AT1G26110), RBP45a (AT5G54900), RBP45b (AT1G11650), RBP45c (AT4G27000), RBP47a (AT1G49600), RBP47b (AT3G19130), UBP1a (AT1G54080), UBP1b (AT1G17370), UBP1c (AT3G14100), G3BP7 (AT5G48650), eIF4a (AT1G54270).

## ACKNOWLEDGEMENTS

We wish to express our gratitude to Dr. Gerardo del Toro-de León for the *ubp1a* seeds, Dr. Julia Bailey-Serres for the *ubp1c* seeds and Dr. Takahiro Hamada for the eIF4A-GFP line.

## AUTHOR CONTRIBUTION

G.H., S.L.G and A.H. designed the experiments and wrote the manuscript. G.H., S.L.G. and A.H. and conducted the experiments analyzed the data. G.H. designed the figures. Polysome analysis was conducted and analyzed together with A.M. and J.H. All authors edited the manuscript and approved the final version.

---

**Supplemental Figure 1:**
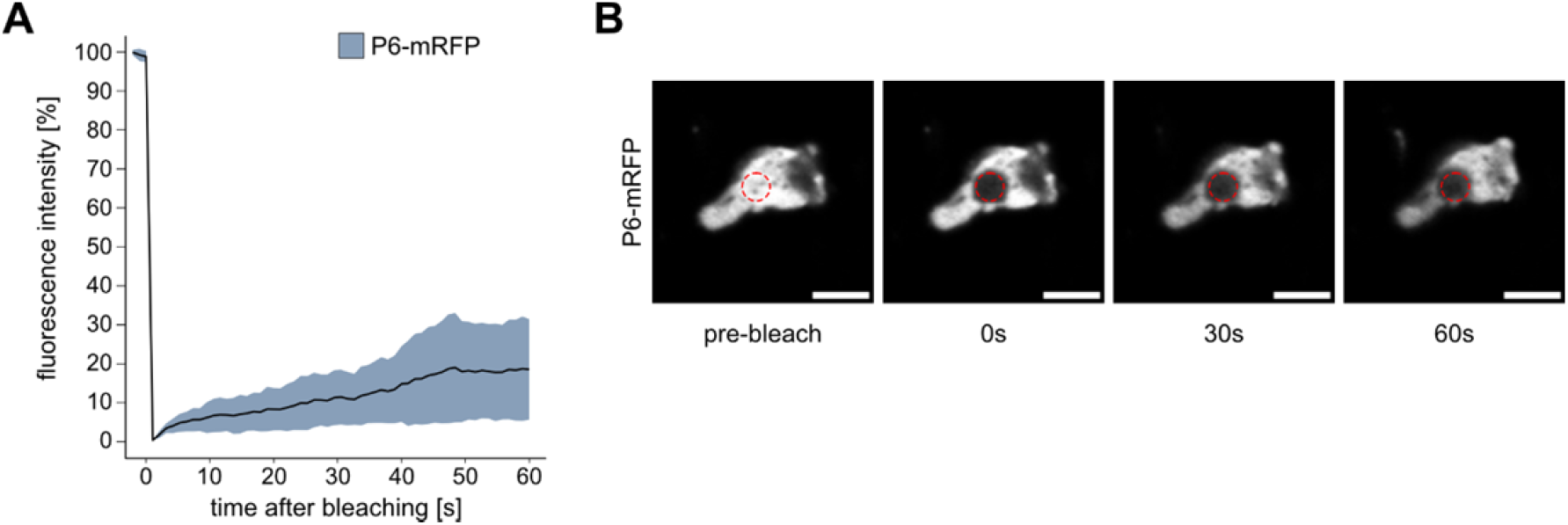
P6-mRFP mobility in viral factories. Supports Figure 1. **(A)** FRAP analysis of P6-mRFP (n=14) in viral factories at 21 dpi, showing normalized fluorescence intensities plotted against time after bleaching. Solid lines represent mean, shades denote ± standard deviation. **(B)** Representative images from FRAP analysis in (A) at indicated time-points. Photobleached region is indicated by a red outline. Scale bars = 5 μm.

**Supplemental Figure 2:**
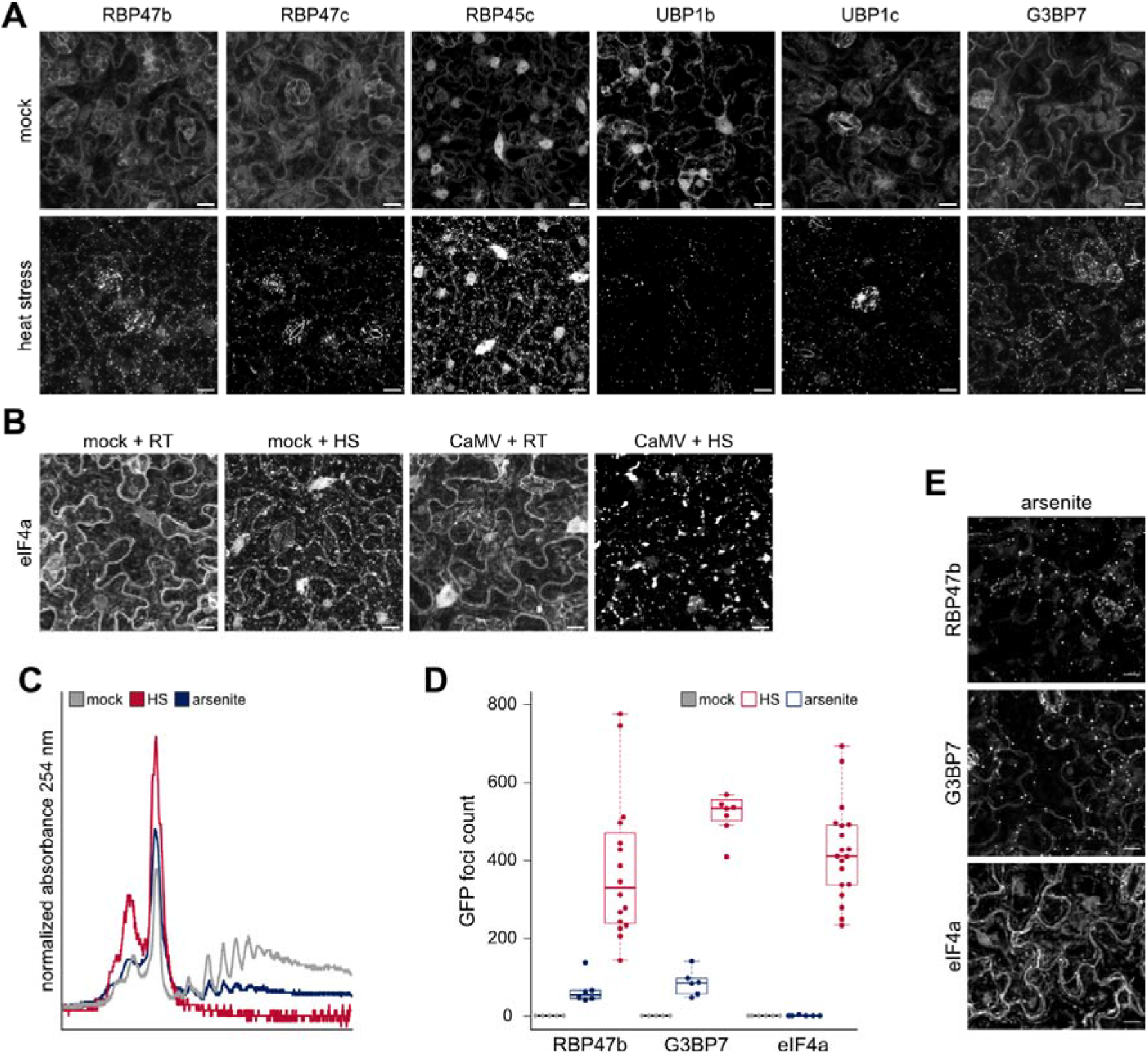
Stress granule proteins localize to viral factories. Supports Figure 2. **(A**) Localization of canonical SG markers in uninfected control conditions (upper panel) and after 30 min of 38 °C heat stress (Lower panel). Representative images are composed of confocal z-stack projections (Scale bars = 10 μm). **(B)** Localization of eIF4A-GFP in control and heat stress conditions. Representative images are composed of confocal z-stack projections (Scale bars = 10 μm). **(C)** Polysome profiles of untreated mock, arsenite or heat stressed GFP-RBP47b plants. **(D)** GFP-foci counts before (mock) and after 1 mM 2h arsenite or 30 min of 38 °C heat treatments for GFP-RBP47b, GFP-G3BP7 and eIF4A-GFP in 100 × 100 μm^2^. Counts were averaged from at least six replicates with a custom ImageJ pipeline. **(B)** Localization of SG markers after 1 mM 2h arsenite treatment. Representative images are composed of confocal z-stack projections (Scale bars = 10 μm).

**Supplemental Figure 3:**
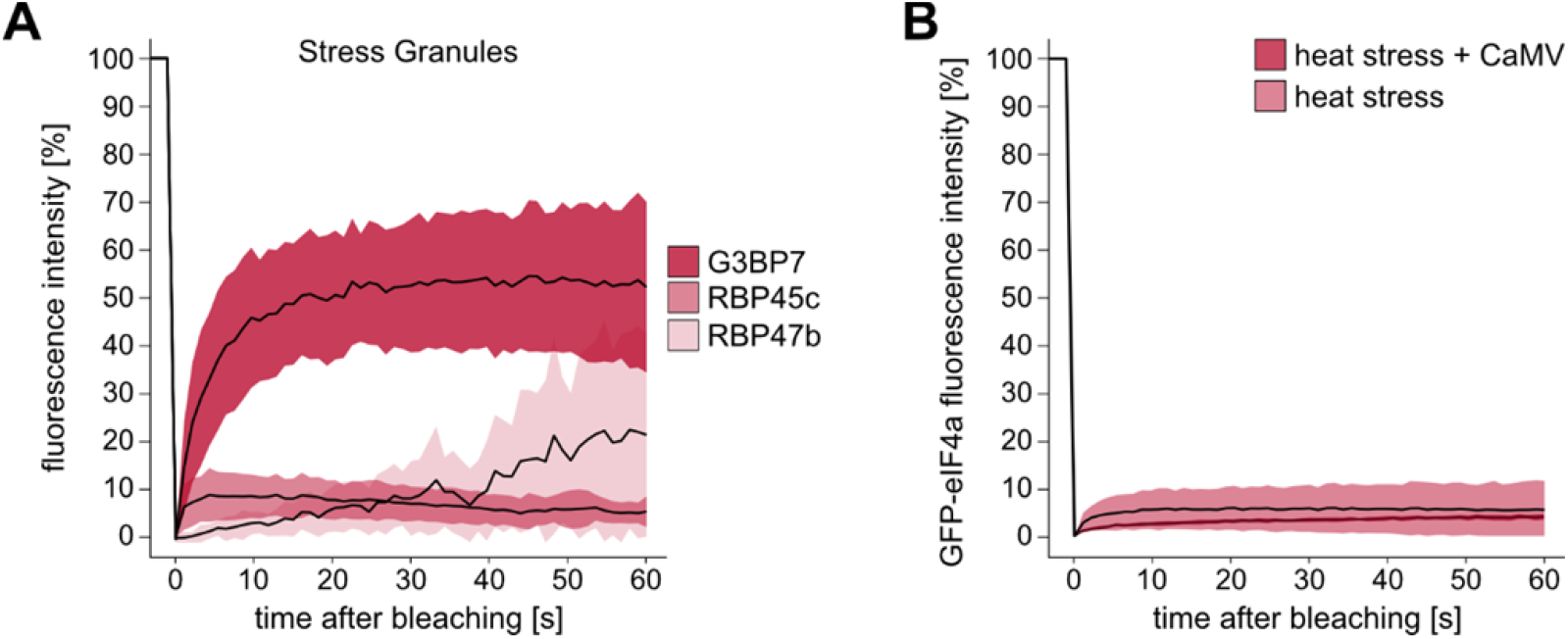
Protein mobility in heat induced SGs. Supports Figure 4. **(A**) Fluorescence recovery of indicated proteins in SG after 30 min of 38 °C in FRAP analysis of uninfected tissue. Normalized fluorescence intensities are plotted against time after bleaching (n=37-41). **(B)** Fluorescence recovery of eIF4A-GFP after photobleaching in VFs (n=7) and SGs (n=60) after 30 min of 38 °C. Normalized fluorescence intensities are plotted against time after bleaching.

**Supplemental Figure 4:**
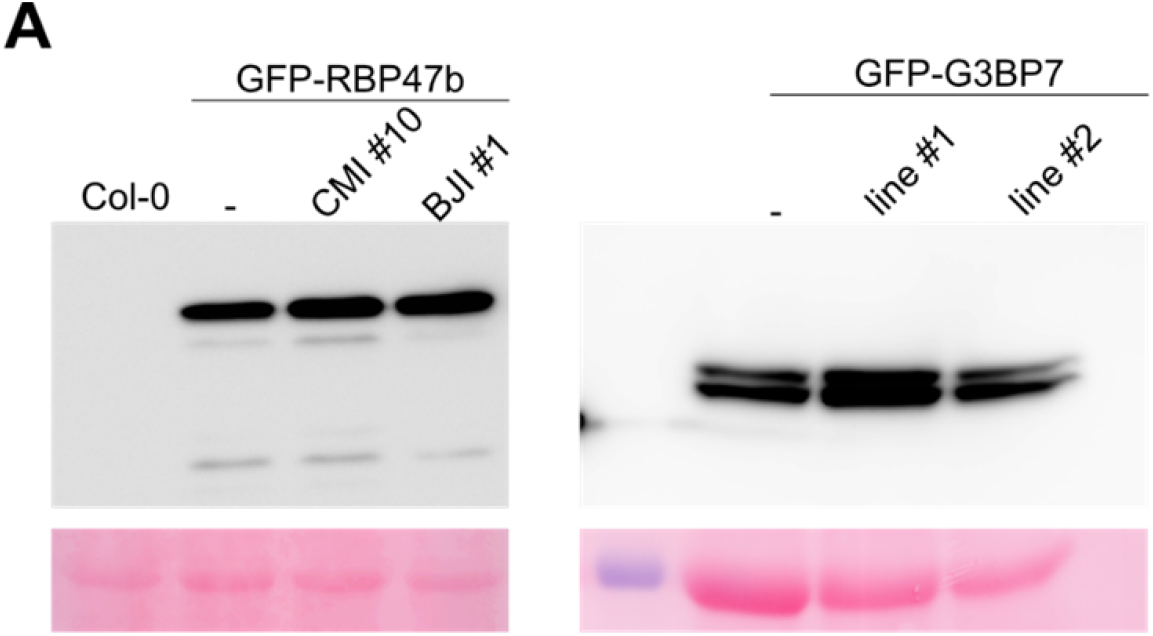
P6 inhibits Stress Granule formation. Supports Figure 6. **(A**) Protein levels of GFP-RBP47b in parental and P6-mRFP lines CMI #10 and BJI #1 and, the same for GFP-G3BP7 parental and P6-mRFP lines #1 and #2. Both were detected using western blot analysis with anti-GFP and, Ponceau S staining is included below to visualize loading.

## Parsed Citations

**Abulfaraj, AA, Mariappan, K., Bigeard, J., Manickam, P., Blilou, I., Guo, X., Al-Babili, S., Pflieger, D., Hirt, H., and Rayapuram, N. (2018). The Arabidopsis homolog of human G3BP1 is a key regulator of stomatal and apoplastic immunity. Life Sci Alliance 1, e201800046.**

Google Scholar: <u>Author Only Title Only Author and Title</u>

**Alberti, S., Gladfelter, A, and Mittag, T. (2019). Considerations and Challenges in Studying Liquid-Liquid Phase Separation and Biomolecular Condensates. Cell 176, 419-434.**

Google Scholar: <u>Author Only Title Only Author and Title</u>

**Alers-Velazquez, R., Jacques, S., Muller, C., Boldt, J., Schoelz, J., and Leisner, S. (2021). Cauliflower mosaic virus P6 inclusion body formation: A dynamic and intricate process. Virology 553, 9-22.**

Google Scholar: <u>Author Only Title Only Author and Title</u>

**Bailey-Serres, J., Sorenson, R., and Juntawong, P. (2009). Getting the message across: cytoplasmic ribonucleoprotein complexes. Trends Plant Sci 14, 443-453.**

Google Scholar: <u>Author Only Title Only Author and Title</u>

**Bak, A, Gargani, D., Macia, J.L., Malouvet, E., Vernerey, M.S., Blanc, S., and Drucker, M. (2013). Virus factories of cauliflower mosaic virus are virion reservoirs that engage actively in vector transmission. Journal of virology 87, 12207-12215.**

Google Scholar: <u>Author Only Title Only Author and Title</u>

**Bernstam, L., and Nriagu, J. (2000). Molecular aspects of arsenic stress. J Toxicol Environ Health B Crit Rev 3, 293-322.**

Google Scholar: <u>Author Only Title Only Author and Title</u>

**Blevins, T., Rajeswaran, R., Aregger, M., Borah, B.K., Schepetilnikov, M., Baerlocher, L., Farinelli, L., Meins, F., Jr., Hohn, T., and Pooggin, M.M. (2011). Massive production of small RNAs from a non-coding region of Cauliflower mosaic virus in plant defense and viral counter-defense. Nucleic acids research 39, 5003-5014.**

Google Scholar: <u>Author Only Title Only Author and Title</u>

**Bolte, S., and Cordelières, F.P. (2006). A guided tour into subcellular colocalization analysis in light microscopy. J Microsc 224, 213-232.**

Google Scholar: <u>Author Only Title Only Author and Title</u>

**Bolte, S., and Cordelieres, F.P. (2006). A guided tour into subcellular colocalization analysis in light microscopy. J Microsc-Oxford 224, 213-232.**

Google Scholar: <u>Author Only Title Only Author and Title</u>

**Boncella, A.E., Shattuck, J.E., Cascarina, S.M., Paul, K.R., Baer, M.H., Fomicheva, A., Lamb, A.K., and Ross, E.D. (2020). Composition-based prediction and rational manipulation of prion-like domain recruitment to stress granules. Proceedings of the National Academy of Sciences of the United States of America 117, 5826-5835.**

Google Scholar: <u>Author Only Title Only Author and Title</u>

**Buchan, J.R., and Parker, R. (2009). Eukaryotic stress granules: the ins and outs of translation. Mol Cell 36, 932-941.**

Google Scholar: <u>Author Only Title Only Author and Title</u>

**Buchan, J.R., Kolaitis, R.M., Taylor, J.P., and Parker, R. (2013). Eukaryotic stress granules are cleared by autophagy and Cdc48/VCP function. Cell 153, 1461-1474.**

Google Scholar: <u>Author Only Title Only Author and Title</u>

**Bureau, M., Leh, V., Haas, M., Geldreich, A, Ryabova, L., Yot, P., and Keller, M. (2004). P6 protein of Cauliflower mosaic virus, a translation reinitiator, interacts with ribosomal protein L13 from Arabidopsis thaliana. The Journal of general virology 85, 3765-3775.**

Google Scholar: <u>Author Only Title Only Author and Title</u>

**Champagne, J., Benhamou, N., and Leclerc, D. (2004). Localization of the N-terminal domain of cauliflower mosaic virus coat protein precursor. Virology 324, 257-262.**

Google Scholar: <u>Author Only Title Only Author and Title</u>

**Chantarachot, T., and Bailey-Serres, J. (2018). Polysomes, Stress Granules, and Processing Bodies: A Dynamic Triumvirate Controlling Cytoplasmic mRNA Fate and Function. Plant physiology 176, 254-269.**

Google Scholar: <u>Author Only Title Only Author and Title</u>

**Clough, S.J., and Bent, A.F. (1998). Floral dip: a simplified method for Agrobacterium -mediated transformation of Arabidopsis thaliana. The Plant Journal 16, 735-743.**

Google Scholar: <u>Author Only Title Only Author and Title</u>

**Conti, G.G., Vegetti, G., Bassi, M., and Favali, M.A (1972). Some ultrastructural and cytochemical observations on Chinese cabbage leaves infected with cauliflower mosaic virus. Virology 47, 694-700.**

Google Scholar: <u>Author Only Title Only Author and Title</u>

**Corbet, G.A., Burke, J.M., and Parker, R. (2021). ADAR1 limits stress granule formation through both translation-dependent and translation-independent mechanisms. J Cell Sci 134.**

Google Scholar: <u>Author Only Title Only Author and Title</u>

**Decker, C.J., and Parker, R. (2012). P-bodies and stress granules: possible roles in the control of translation and mRNA degradation. Cold Spring Harb Perspect Biol 4, a012286.**

Google Scholar: <u>Author Only Title Only Author and Title</u>

**Dember, L.M., Kim, N.D., Liu, K.Q., and Anderson, P. (1996). Individual RNA recognition motifs of TIAr1 and TIAR have different RNA binding specificities. J Biol Chem 271, 2783-2788.**

Google Scholar: <u>Author Only Title Only Author and Title</u>

**Emara, M.M., and Brinton, M.A (2007). Interaction of TIA-1/TIAR with West Nile and dengue virus products in infected cells interferes with stress granule formation and processing body assembly. Proceedings of the National Academy of Sciences of the United States of America 104, 9041-9046.**

Google Scholar: <u>Author Only Title Only Author and Title</u>

**Fare, C.M., Villani, A, Drake, L.E., and Shorter, J. (2021). Higher-order organization of biomolecular condensates. Open Biol 11, 210137.**

Google Scholar: <u>Author Only Title Only Author and Title</u>

**Frydryskova, K., Masek, T., and Pospisek, M. (2020). Changing faces of stress: Impact of heat and arsenite treatment on the composition of stress granules. Wiley Interdiscip Rev RNA 11, e1596.**

Google Scholar: <u>Author Only Title Only Author and Title</u>

**Grefen, C., Donald, N., Hashimoto, K., Kudla, J., Schumacher, K., and Blatt, M.R. (2010). A ubiquitin-10 promoter-based vector set for fluorescent protein tagging facilitates temporal stability and native protein distribution in transient and stable expression studies. The Plant Journal 64, 355-365.**

Google Scholar: <u>Author Only Title Only Author and Title</u>

**Gutierrez-Beltran, E., Moschou, P.N., Smertenko, AP., and Bozhkov, P.V. (2015). Tudor staphylococcal nuclease links formation of stress granules and processing bodies with mRNA catabolism in Arabidopsis. The Plant cell 27, 926-943.**

Google Scholar: <u>Author Only Title Only Author and Title</u>

**Gutierrez-Beltran, E., Elander, P.H., Dalman, K., Dayhoff, G.W., 2nd, Moschou, P.N., Uversky, V.N., Crespo, J.L., and Bozhkov, P.V. (2021). Tudor staphylococcal nuclease is a docking platform for stress granule components and is essential for SnRK1 activation in Arabidopsis. The EMBO journal 40, e105043.**

Google Scholar: <u>Author Only Title Only Author and Title</u>

**Guzikowski, AR., Chen, Y.S., and Zid, B.M. (2019). Stress-induced mRNP granules: Form and function of processing bodies and stress granules. Wiley Interdiscip Rev RNA 10, e1524.**

Google Scholar: <u>Author Only Title Only Author and Title</u>

**Haas, G., Azevedo, J., Moissiard, G., Geldreich, A., Himber, C., Bureau, M., Fukuhara, T., Keller, M., and Voinnet, O. (2008). Nuclear import of CaMV P6 is required for infection and suppression of the RNA silencing factor DRB4. The EMBO journal 27, 2102-2112.**

Google Scholar: <u>Author Only Title Only Author and Title</u>

**Hafren, A, Lohmus, A, and Makinen, K. (2015). Formation of Potato Virus A-Induced RNA Granules and Viral Translation Are Interrelated Processes Required for Optimal Virus Accumulation. PLoS pathogens 11, e1005314.**

Google Scholar: <u>Author Only Title Only Author and Title</u>

**Hafren, A, Macia, J.L., Love, A.J., Milner, J.J., Drucker, M., and Hofius, D. (2017). Selective autophagy limits cauliflower mosaic virus infection by NBR1-mediated targeting of viral capsid protein and particles. Proceedings of the National Academy of Sciences of the United States of America 114, E2026-E2035.**

Google Scholar: <u>Author Only Title Only Author and Title</u>

**Hamada, T., Yako, M., Minegishi, M., Sato, M., Kamei, Y., Yanagawa, Y., Toyooka, K., Watanabe, Y., and Hara-Nishimura, I. (2018). Stress granule formation is induced by a threshold temperature rather than a temperature difference in Arabidopsis. J Cell Sci 131.**

Google Scholar: <u>Author Only Title Only Author and Title</u>

**Hapiak, M., Li, Y., Agama, K., Swade, S., Okenka, G., Falk, J., Khandekar, S., Raikhy, G., Anderson, A., Pollock, J., Zellner, W., Schoelz, J., and Leisner, S.M. (2008). Cauliflower mosaic virus gene VI product N-terminus contains regions involved in resistance-breakage, self-association and interactions with movement protein. Virus Res 138, 119-129.**

Google Scholar: <u>Author Only Title Only Author and Title</u>

**Harries, P.A., Palanichelvam, K., Yu, W., Schoelz, J.E., and Nelson, R.S. (2009). The cauliflower mosaic virus protein P6 forms motile inclusions that traffic along actin microfilaments and stabilize microtubules. Plant physiology 149, 1005-1016.**

Google Scholar: <u>Author Only Title Only Author and Title</u>

**Himmelbach, A, Chapdelaine, Y., and Hohn, T. (1996). Interaction between cauliflower mosaic virus inclusion body protein and capsid protein: implications for viral assembly. Virology 217, 147-157.**

Google Scholar: <u>Author Only Title Only Author and Title</u>

**Hoffmann, G., Mahboubi, A, Bente, H., Garcia, D., Hanson, J., and Hafr, N.A (2022). Arabidopsis RNA processing body components LSM1 and DCP5 aid in the evasion of translational repression during Cauliflower mosaic virus infection. The Plant cell 34, 3128-3147.**

Google Scholar: <u>Author Only Title Only Author and Title</u>

**Hohn, T. (2015). RNA based viral silencing suppression in plant pararetroviruses. Front Plant Sci 6, 398.**

Google Scholar: <u>Author Only Title Only Author and Title</u>

**Jaafar, Z.A., and Kieft, J.S. (2019). Viral RNA structure-based strategies to manipulate translation. Nature reviews. Microbiology 17, 110-123.**

Google Scholar: <u>Author Only Title Only Author and Title</u>

**Kedersha, N., and Anderson, P. (2007). Mammalian stress granules and processing bodies. Methods Enzymol 431, 61-81.**

Google Scholar: <u>Author Only Title Only Author and Title</u>

**Kedersha, N., Cho, M.R., Li, W., Yacono, P.W., Chen, S., Gilks, N., Golan, D.E., and Anderson, P. (2000). Dynamic shuttling of TIA-1 accompanies the recruitment of mRNA to mammalian stress granules. The Journal of cell biology 151, 1257-1268.**

Google Scholar: <u>Author Only Title Only Author and Title</u>

**Kedersha, N., Stoecklin, G., Ayodele, M., Yacono, P., Lykke-Andersen, J., Fritzler, M.J., Scheuner, D., Kaufman, R.J., Golan, D.E., and Anderson, P. (2005). Stress granules and processing bodies are dynamically linked sites of mRNP remodeling. The Journal of cell biology 169, 871-884.**

Google Scholar: <u>Author Only Title Only Author and Title</u>

**Khong, A, Matheny, T., Jain, S., Mitchell, S.F., Wheeler, J.R., and Parker, R. (2017). The Stress Granule Transcriptome Reveals Principles of mRNA Accumulation in Stress Granules. Mol Cell 68, 808-820 e805.**

Google Scholar: <u>Author Only Title Only Author and Title</u>

**Kitajima, E.W., Lauritis, J.A, and Swift, H. (1969). Fine structure of zinnial leaf tissues infected with dahlia mosaic virus. Virology 39, 240-249.**

Google Scholar: <u>Author Only Title Only Author and Title</u>

**Krapp, S., Greiner, E., Amin, B., Sonnewald, U., and Krenz, B. (2017). The stress granule component G3BP is a novel interaction partner for the nuclear shuttle proteins of the nanovirus pea necrotic yellow dwarf virus and geminivirus abutilon mosaic virus. Virus Res 227, 6-14.**

Google Scholar: <u>Author Only Title Only Author and Title</u>

**Laird, J., McInally, C., Carr, C., Doddiah, S., Yates, G., Chrysanthou, E., Khattab, A., Love, A.J., Geri, C., Sadanandom, A., Smith, B.O., Kobayashi, K., and Milner, J.J. (2013). Identification of the domains of cauliflower mosaic virus protein P6 responsible for suppression of RNA silencing and salicylic acid signalling. The Journal of general virology 94, 2777-2789.**

Google Scholar: <u>Author Only Title Only Author and Title</u>

**Leh, V., Yot, P., and Keller, M. (2000). The cauliflower mosaic virus translational transactivator interacts with the 60S ribosomal subunit protein L18 of Arabidopsis thaliana. Virology 266, 1-7.**

Google Scholar: <u>Author Only Title Only Author and Title</u>

**Lloyd, R.E. (2016). Enterovirus Control of Translation and RNA Granule Stress Responses. Viruses 8, 93.**

Google Scholar: <u>Author Only Title Only Author and Title</u>

**Lorkovic, Z.J., and Barta, *A.* (2002). Genome analysis: RNA recognition motif (RRM) and K homology (KH) domain RNA-binding proteins from the flowering plant Arabidopsis thaliana. Nucleic acids research 30, 623-635.**

Google Scholar: <u>Author Only Title Only Author and Title</u>

**Love, AJ., Geri, C., Laird, J., Carr, C., Yun, B.W., Loake, G.J., Tada, Y., Sadanandom, A, and Milner, J.J. (2012). Cauliflower mosaic virus protein P6 inhibits signaling responses to salicylic acid and regulates innate immunity. PloS one 7, e47535.**

Google Scholar: <u>Author Only Title Only Author and Title</u>

**Lukhovitskaya, N., and Ryabova, L.A (2019). Cauliflower mosaic virus transactivator protein (TAV) can suppress nonsense-mediated decay by targeting VARICOSE, a scaffold protein of the decapping complex. Sci Rep 9, 7042.**

Google Scholar: <u>Author Only Title Only Author and Title</u>

**Lutz, L., Raikhy, G., and Leisner, S.M. (2012). Cauliflower mosaic virus major inclusion body protein interacts with the aphid transmission factor, the virion-associated protein, and gene VII product. Virus Res 170, 150-153.**

Google Scholar: <u>Author Only Title Only Author and Title</u>

**Makinen, K., Lohmus, A, and Pollari, M. (2017). Plant RNA Regulatory Network and RNA Granules in Virus Infection. Front Plant Sci 8, 2093.**

Google Scholar: <u>Author Only Title Only Author and Title</u>

**Martelli, G.P., and Castellano, M.A. (1971). Light and electron microscopy of the intracellular inclusions of cauliflower mosaic virus. The Journal of general virology 13, 133-140.**

Google Scholar: <u>Author Only Title Only Author and Title</u>

**McCue, AD., Nuthikattu, S., Reeder, S.H., and Slotkin, R.K. (2012). Gene expression and stress response mediated by the epigenetic regulation of a transposable element small RNA PLoS genetics 8, e1002474.**

Google Scholar: <u>Author Only Title Only Author and Title</u>

**Miras, M., Miller, W.A., Truniger, V., and Aranda, M.A. (2017). Non-canonical Translation in Plant RNA Viruses. Front Plant Sci 8, 494.**

Google Scholar: <u>Author Only Title Only Author and Title</u>

**Moore, H.M., Bai, B., Boisvert, F.M., Latonen, L., Rantanen, V., Simpson, J.C., Pepperkok, R., Lamond, A.I., and Laiho, M. (2011). Quantitative proteomics and dynamic imaging of the nucleolus reveal distinct responses to UV and ionizing radiation. Mol Cell Proteomics 10, M111 009241.**

Google Scholar: <u>Author Only Title Only Author and Title</u>

**Nakagawa, T., Suzuki, T., Murata, S., Nakamura, S., Hino, T., Maeo, K., Tabata, R., Kawai, T., Tanaka, K., Niwa, Y., Watanabe, Y., Nakamura, K., Kimura, T., and Ishiguro, S. (2007). Improved Gateway Binary Vectors: High-Performance Vectors for Creation of Fusion Constructs in Transgenic Analysis of Plants. Bioscience, Biotechnology, and Biochemistry 71, 2095-2100.**

Google Scholar: <u>Author Only Title Only Author and Title</u>

**Panas, M.D., Varjak, M., Lulla, A, Eng, K.E., Merits, A, Karlsson Hedestam, G.B., and McInerney, G.M. (2012). Sequestration of G3BP coupled with efficient translation inhibits stress granules in Semliki Forest virus infection. Mol Biol Cell 23, 4701-4712.**

Google Scholar: <u>Author Only Title Only Author and Title</u>

**Park, H.S., Himmelbach, A, Browning, K.S., Hohn, T., and Ryabova, L.A. (2001). A plant viral “reinitiation” factor interacts with the host translational machinery. Cell 106, 723-733.**

Google Scholar: <u>Author Only Title Only Author and Title</u>

**Poblete-Duran, N., Prades-Perez, Y., Vera-Otarola, J., Soto-Rifo, R., and Valiente-Echeverria, F. (2016). Who Regulates Whom? An Overview of RNA Granules and Viral Infections. Viruses 8.**

Google Scholar: <u>Author Only Title Only Author and Title</u>

**Pooggin, M.M., and Ryabova, L.A. (2018). Ribosome Shunting, Polycistronic Translation, and Evasion of Antiviral Defenses in Plant Pararetroviruses and Beyond. Front Microbiol 9, 644.**

Google Scholar: <u>Author Only Title Only Author and Title</u>

**Powers, S.K., Holehouse, A.S., Korasick, D.A., Schreiber, K.H., Clark, N.M., Jing, H., Emenecker, R., Han, S., Tycksen, E., Hwang, I., Sozzani, R., Jez, J.M., Pappu, R.V., and Strader, L.C. (2019). Nucleo-cytoplasmic Partitioning of ARF Proteins Controls Auxin Responses in Arabidopsis thaliana. Mol Cell 76, 177-190 e175.**

Google Scholar: <u>Author Only Title Only Author and Title</u>

**Reuper, H., Amari, K., and Krenz, B. (2021). Analyzing the G3BP-like gene family of Arabidopsis thaliana in early turnip mosaic virus infection. Sci Rep 11, 2187.**

Google Scholar: <u>Author Only Title Only Author and Title</u>

**Riggs, C.L., Kedersha, N., Ivanov, P., and Anderson, P. (2020). Mammalian stress granules and P bodies at a glance. J Cell Sci 133.**

Google Scholar: <u>Author Only Title Only Author and Title</u>

**Schepetilnikov, M., Kobayashi, K., Geldreich, A, Caranta, C., Robaglia, C., Keller, M., and Ryabova, L.A. (2011). Viral factor TAV recruits TOR/S6K1 signalling to activate reinitiation after long ORF translation. The EMBO journal 30, 1343-1356.**

Google Scholar: <u>Author Only Title Only Author and Title</u>

**Schoelz, J.E., and Leisner, S. (2017). Setting Up Shop: The Formation and Function of the Viral Factories of Cauliflower mosaic virus. Front Plant Sci 8, 1832.**

Google Scholar: <u>Author Only Title Only Author and Title</u>

**Schoelz, J.E., Goldberg, K.B., and Kiernan, J. (1991). Expression of Cauliflower Mosaic-Virus (Camv) Gene-Vi in Transgenic Nicotiana-Bigelovii Complements a Strain of Camv Defective in Long-Distance Movement in Nontransformed N-Bigelovii. Mol Plant Microbe In 4, 350-355.**

Google Scholar: <u>Author Only Title Only Author and Title</u>

**Sfakianos, A.P., Mellor, L.E., Pang, Y.F., Kritsiligkou, P., Needs, H., Abou-Hamdan, H., Desaubry, L., Poulin, G.B., Ashe, M.P., and Whitmarsh, AJ.** (2018). The mTOR-S6 kinase pathway promotes stress granule assembly. Cell death and differentiation **25**, 1766–1780.

Google Scholar: <u>Author Only Title Only Author and Title</u>

**Shepherd, R.J. (1976). DNA viruses of higher plants. Advances in virus research 20, 305-339.**

Google Scholar: <u>Author Only Title Only Author and Title</u>

**Shukla, A, Hoffmann, G., Kushwaha, N.K., Lopez-Gonzalez, S., Hofius, D., and Hafren, A. (2021). Salicylic acid and the viral virulence factor 2b regulate the divergent roles of autophagy during cucumber mosaic virus infection. Autophagy, 1-13.**

Google Scholar: <u>Author Only Title Only Author and Title</u>

**Sorenson, R., and Bailey-Serres, J. (2014). Selective mRNA sequestration by OLIGOURIDYLATE-BINDING PROTEIN 1 contributes to translational control during hypoxia in Arabidopsis. Proceedings of the National Academy of Sciences of the United States of America 111, 2373-2378.**

Google Scholar: <u>Author Only Title Only Author and Title</u>

**Souquere, S., Mollet, S., Kress, M., Dautry, F., Pierron, G., and Weil, D. (2009). Unravelling the ultrastructure of stress granules and associated P-bodies in human cells. J Cell Sci 122, 3619-3626.**

Google Scholar: <u>Author Only Title Only Author and Title</u>

**Spector, D.L. (2006). SnapShot: Cellular bodies. Cell 127, 1071.**

Google Scholar: <u>Author Only Title Only Author and Title</u>

**Stern-Ginossar, N., Thompson, S.R., Mathews, M.B., and Mohr, I. (2019). Translational Control in Virus-Infected Cells. Cold Spring Harb Perspect Biol 11.**

Google Scholar: <u>Author Only Title Only Author and Title</u>

**Teixeira, D., Sheth, U., Valencia-Sanchez, M.A., Brengues, M., and Parker, R. (2005). Processing bodies require RNA for assembly and contain nontranslating mRNAs. Rna 11, 371-382.**

Google Scholar: <u>Author Only Title Only Author and Title</u>

**Torres-Barcelo, C., Martin, S., Daros, J.A., and Elena, S.F. (2008). From hypo- to hypersuppression: effect of amino acid substitutions on the RNA-silencing suppressor activity of the Tobacco etch potyvirus HC-Pro. Genetics 180, 1039-1049.**

Google Scholar: <u>Author Only Title Only Author and Title</u>

**Ustun, S., Hafren, A, Liu, Q., Marshall, R.S., Minina, E.A., Bozhkov, P.V., Vierstra, R.D., and Hofius, D. (2018). Bacteria Exploit Autophagy for Proteasome Degradation and Enhanced Virulence in Plants. The Plant cell 30, 668-685.**

Google Scholar: <u>Author Only Title Only Author and Title</u>

**Uversky, V.N. (2017). Intrinsically disordered proteins in overcrowded milieu: Membrane-less organelles, phase separation, and intrinsic disorder. Curr Opin Struct Biol 44, 18-30.**

Google Scholar: <u>Author Only Title Only Author and Title</u>

**Weber, C., Nover, L., and Fauth, M. (2008). Plant stress granules and mRNA processing bodies are distinct from heat stress granules. The Plant journal: for cell and molecular biology 56, 517-530.**

Google Scholar: <u>Author Only Title Only Author and Title</u>

**Yang, P., Mathieu, C., Kolaitis, R.M., Zhang, P., Messing, J., Yurtsever, U., Yang, Z., Wu, J., Li, Y., Pan, Q., Yu, J., Martin, E.W., Mittag, T., Kim, H.J., and Taylor, J.P. (2020). G3BP1 Is a Tunable Switch that Triggers Phase Separation to Assemble Stress Granules. Cell 181, 325-345 e328.**

Google Scholar: <u>Author Only Title Only Author and Title</u>

**Youn, J.Y., Dyakov, B.J.A, Zhang, J., Knight, J.D.R., Vernon, R.M., Forman-Kay, J.D., and Gingras, AC. (2019). Properties of Stress Granule and P-Body Proteomes. Mol Cell 76, 286-294.**

Google Scholar: <u>Author Only Title Only Author and Title</u>

**Youn, J.Y., Dunham, W.H., Hong, S.J., Knight, J.D.R., Bashkurov, M., Chen, G.I., Bagci, H., Rathod, B., MacLeod, G., Eng, S.W.M., Angers, S., Morris, Q., Fabian, M., Cote, J.F., and Gingras, AC. (2018). High-Density Proximity Mapping Reveals the Subcellular Organization of mRNA-Associated Granules and Bodies. Mol Cell 69, 517-532 e511.**

Google Scholar: <u>Author Only Title Only Author and Title</u>

**Zhang, X.P., Liu, D.S., Yan, T., Fang, X.D., Dong, K., Xu, J., Wang, Y., Yu, J.L., and Wang, X.B. (2017). Cucumber mosaic virus coat protein modulates the accumulation of 2b protein and antiviral silencing that causes symptom recovery in planta. PLoS pathogens 13, e1006522.**

Google Scholar: <u>Author Only Title Only Author and Title</u>

**Zvereva, A.S., Golyaev, V., Turco, S., Gubaeva, E.G., Rajeswaran, R., Schepetilnikov, M.V., Srour, O., Ryabova, L.A., Boller, T., and Pooggin, M.M. (2016). Viral protein suppresses oxidative burst and salicylic acid-dependent autophagy and facilitates bacterial growth on virus-infected plants. The New phytologist.**

Google Scholar: <u>Author Only Title Only Author and Title</u>

